# Two decades of satellite images reveal the spatial and temporal dynamics of leafy spurge invasion and improve species distribution models

**DOI:** 10.1101/2025.05.09.653185

**Authors:** Thomas A. Lake, Ryan D. Briscoe Runquist, David A. Moeller

**Affiliations:** University of Minnesota, Plant and Microbial Biology Department, 140 Gortner Laboratory, 1479 Gortner Ave, St. Paul, MN 55108; North Carolina State University, Center for Geospatial Analytics, 5112 Jordan Hall, 2800 Faucette Dr, Raleigh, NC 27695

**Author notes:** Co-first authors (Thomas A. Lake, Ryan D. Briscoe Runquist).

**Keywords:** process-based distribution models, population dynamics, convolutional neural network, deep learning, biological invasions, satellite image time-series, Landsat satellite imagery

## Abstract

Developing accurate and cost-effective methods to detect and predict the spread of invasive species remains an ongoing challenge. Species distribution models (SDMs) are used to predict invasion under current and future climates. However, occurrence datasets are often spatially and temporally biased because they are collected in an unstructured and opportunistic manner. Remote sensing individual species over broad spatial and temporal scales has become increasingly feasible with the accumulation of satellite images and the development of convolutional neural networks. In this study, we used a 21-year archival Landsat satellite imagery (2000-2020) to train a temporal convolutional neural network model to predict the probability of occurrence of the invasive species, leafy spurge (*Euphorbia virgata*), across Minnesota. We validated our predictions with an independent dataset. First, we show that that leafy spurge has expanded from 1,067 km² occupied in 2000–2002 to 7,156 km² in 2018-2020, a 570% increase. Surprisingly, drought severity modulated the predicted area invaded and was associated with fluctuations over the study period. Second, we tracked changes in probability over time for individual pixels and showed that invasion has been concentrated in two largely disjunct regions of Minnesota. Third, our remotely-sensed occurrence dataset and community science dataset were biased to roadsides, although the latter was more severely biased. Last, we showed that SDMs built using remotely-sensed occurrences had higher discrimination, were less overfit, and had higher performance outside of urban areas. Overall, twenty one years of archival satellite imagery provided valuable insight into the spatial and temporal population dynamics of leafy spurge invasion and improved forecasts of future invasion by reducing spatial bias.

## INTRODUCTION

Unbiased assessment of the distribution and abundance of individual species is fundamental to basic and applied problems in ecology, biogeography, and conservation (Meyer et al., 2015). However, occurrence datasets are typically spatially biased, which has compromised our ability to fairly test biogeographic predictions (e.g. abundant center hypothesis) and to accurately forecast species’ range shifts (Garcia-Rosello et al., 2023; Hughes et al., 2021). Recent developments in remote sensing have provided avenues for identifying individual species from satellite images with considerable accuracy (Randin et al., 2020; Wang & Gamon, 2019). Although rarely attempted, remote sensing models increasingly have the potential to characterize the unbiased distribution and population dynamics of individual species over large geographic areas and long time periods (He et al., 2015; Randin et al., 2020).

Among the most frequent and important uses of occurrence datasets is for the construction of species distribution models or environmental niche models (hereafter SDMs) (Elith & Leathwick, 2009; Guisan & Zimmermann, 2000; Miller, 2010). These models are widely used to forecast range shifts with climate change, anticipate invasions, and identify habitat for conservation (Franklin, 2023). In most models, occurrences are acquired from publicly available databases that accumulate records from many sources and over long time periods. For example, records in the Global Biodiversity Information Facility (GBIF) are collated from biodiversity inventories, targeted surveys (e.g. documenting invasive species in a particular jurisdiction), observations by scientists, and observations by community scientists (Di Cecco et al., 2021; Larson et al., 2020). Although these occurrence datasets are often large, they rarely sample the landscape randomly or accurately represent changes in abundance over time.

Similarly, areas devoid of records may not represent true absences if sampling is incomplete in remote areas. Overall, sampling biases have been found across taxonomic categories, sampling methods, and even in the largest datasets (Beck et al., 2014; Courter et al., 2013; Di Cecco et al., 2021; Geurts et al., 2023; Meyer et al., 2016; Santos et al., 2020; Shirey et al., 2021).

Spatial bias in occurrence datasets is problematic because it influences our understanding of the spatial dynamics of invasion and the distribution of occurrence-environment relationships. When species’ distributions are rapidly changing (e.g. invasions), occurrences may be concentrated in regions first colonized and poorly represented at expanding range margins.

Similarly, observations may be concentrated spatially or temporally where and when effort is greater, such as in urban centers or during times of increased awareness and participation in community science (Bowler et al., 2022). When data collection biases translate to biases in environmental associations, species distribution models often fail to accurately predict suitable but unoccupied areas (Briscoe Runquist et al., 2019; Lustenhouwer & Parker, 2022; Phillips et al., 2009; Yates et al., 2018). This problem is particularly pronounced at expanding range margins where observations of occurrence are often less common by virtue of difference in prevalence and sampling effort (Bowler et al., 2022; Isaac & Pocock, 2015). Considerable effort has been dedicated to developing methods that reduce spatial bias (Franklin, 2023). While those approaches are helpful in some cases, they typically do not eliminate bias and generating accurate forecasts of range expansion continues to be challenging (Fourcade et al., 2014; Jarnevich et al., 2015; Lake et al., 2020; Syfert et al., 2013).

Traditional SDMs are built based on patterns of occurrence but do not account for variation in abundance or demography. Process-based SDMs relate biological processes (such as demographic factors) to environmental variability (Briscoe et al., 2019; Dormann et al., 2012; Higgins et al., 2020). Process-based models may better predict habitat suitability and the extent of range shifts with climate change (Dormann et al., 2012; Van der Meersch et al., 2024). For instance, incorporating population growth rates can help differentiate environments with expanding, stable, or declining populations (Eckhart et al., 2011; Yu et al., 2020). Despite their advantages, these models are rare because they require long-term monitoring (or experimentation) across a species’ range (Briscoe et al., 2019; Dormann et al., 2012).

Advances in remote sensing, satellite imagery systems, and computing have enabled detailed mapping of plant populations (Bradley, 2014; Cerrejón et al., 2021; Müllerová et al., 2023). Multispectral imagery archives are a rapidly expanding resource that can be used to map all levels of biodiversity across the globe (Randin et al 2020). The Landsat satellite imagery archive is unique due to its deep temporal catalogue: the global dataset includes images captured approximately every 8-16 days for over 50 years (1972-present). To date, the dataset has been employed for long-term plant community and ecosystem monitoring projects (Bradley & Mustard, 2006; Singh & Glenn, 2009; Wulder et al., 2016, 2022). In addition, shifts in distribution and abundance have been detected by comparing historical and contemporary imagery. For example, Landsat imagery has revealed the expansion of plant communities dominated by exotic grasses to higher elevations over three decades, aligning with climate change projections (Smith et al., 2022). Remote sensing individual species using a time-series of images has the potential to infer demographic parameters and inform process-based SDMs to better forecast range shifts.

Biological invasions threaten native biodiversity, disrupt ecosystem function, and burden economies worldwide (Fantle-Lepczyk et al., 2022; Pyšek et al., 2020; Seebens et al., 2017). Given these impacts, there is a need to improve forecasts of invasive species’ potential distributions to inform management strategies (Hanley & Roberts, 2019; Pyšek et al., 2020; Seebens et al., 2020). Recently, remote sensing approaches have accurately detected populations of individual invasive plant species (Asner et al., 2008; Carter et al., 2009; Chen et al., 2020; Clinton et al., 2010; Dahal et al., 2022; Hunt & Parker Williams, 2006; Lake et al., 2022; Mattilio et al., 2023; Ren et al., 2021). Imagery time series are likely of particular value for detecting occurrences in remote, unsampled regions and for determining regions where new invasion is occurring. In the context of management, remotely-sensed occurrences can be leveraged to overcome spatial biases in community science observations and build more accurate forecasts for range shifts with climate change (He et al., 2015).

In this study, we used a 21-year dataset of Landsat imagery for Minnesota, USA to quantify changes in the spatial distribution and population dynamics of leafy spurge (*Euphorbia virgata*; Euphorbiaceae; Levin & Gillespie, 2017), among the most economically damaging invasive plants in the US (Fantle-Lepczyk et al., 2022). To detect leafy spurge from imagery, we trained a temporal convolutional neural network, used a fully withheld testing dataset to evaluate classification accuracy, and interrogated the circumstances of classification failures. We then calculated population demographic changes over 21 years for each pixel and estimated the area of invasion across Minnesota through time. We also examined whether climatic variation during this period influenced detection and inferences about the extent of invasion. Last, we compared the performance of species distribution models based on (1) remotely-sensed occurrences, (2) remotely-sensed inferences of population dynamics, and (3) a community science dataset of occurrences.

## MATERIALS AND METHODS

### Study system

Leafy spurge is an herbaceous perennial plant that was introduced to North America from Eurasia in the 1800s. It has spread rapidly since introduction, approximately doubling its invaded range every decade through the early 2000s (Leistritz et al., 1992; Leitch et al., 1994). It now occupies nearly two million hectares of grasslands, rangelands, and open forest edges in the northern United States and southern Canada (DiTomaso, 2000; Duncan et al., 2004; Dunn, 1979; Leistritz et al., 1992) and is among the most economically damaging invasive plants in the US with losses between 1960-2020 totaling over $1 billion (Bangsund et al., 1999; DiTomaso, 2000; Duncan et al., 2004; Fantle-Lepczyk et al., 2022; Leitch et al., 1994). Plants emerge from dormancy in early spring before most herbaceous plants and flower in late spring (Bangsund et al., 1999; Lym, 1998). The flowers are subtended by bracts that have a distinct yellow-green color. When damaged, plants produce a latex that can be toxic to livestock, which in turn affects the value of invaded rangelands (Messersmith et al., 1985).

Prior remote sensing studies have shown that the distinct spectral signal of its flowers is particularly important for distinguishing leafy spurge from co-occurring species (Casady et al., 2005; Hunt et al., 2004; Lake et al., 2022; Mattilio et al., 2023; Stitt et al., 2006). Recently, the use of a temporal series of images has shown that the phenology of emergence, flowering, and senescence provides an important signature that facilitates detection even with lower resolution imagery (Lake et al., 2022).

Leafy spurge was introduced to southwestern Minnesota in ca. 1890 and now occurs across a substantial fraction of Minnesota, USA (225,000 square km). By the 1930s, it had spread across the grasslands but occurrences in deciduous and boreal forests were uncommon (Hanson & Rudd, 1933). Leafy spurge remained rare in boreal forests until the 1990s.

### Developing and evaluating a remote sensing model

#### Leafy spurge occurrence data

We downloaded community science occurrences between 2000 and 2020 from the Early Detection and Distribution Mapping System (EDDMapS.org) and iNaturalist (iNaturalist.org) for Minnesota (Table S1). We filtered occurrences by removing duplicate, imprecise (less than 0.01 decimal degrees of precision), and implausible (e.g., over water) records. We kept only "verified" occurrences from EDDMapS and "research grade" occurrences from iNaturalist. To reduce spatial autocorrelation among occurrences, we applied spatial point thinning to preserve one point per 100 meters. In total, we obtained 3,345 occurrences for use in downstream analyses.

#### Landsat satellite imagery

We accessed Landsat 5 TM, 7 ETM+, and 8 OLI surface reflectance products (2000-2020) via Google Earth Engine (Gorelick et al., 2017). Landsat images provide a moderate resolution (30 meter) and 16-day revisit period that is ideal for tracking vegetation dynamics (Vogelmann et al., 2016). We used data from the visible, near infrared, and shortwave bands in addition to the normalized difference vegetation index [NDVI = (near infrared - red) / (near infrared + red)], which is a proxy for seasonal photosynthetic activity (Gamon et al., 1995). We first processed imagery by removing pixels with cloud, cloud shadow, and radiometric saturation using quality assessment bitmasks. We then created three seasonal windows per year, each of which represents a two-month period: March-April, May-June, and July-August. These windows correspond to the timing of key events in leafy spurge’s growing season: emergence, flowering, and senescence. For each seasonal window, we calculated the median pixel value for each spectral band for all images within that two-month window (additional methods for image processing can be found in Supplementary Methods 1.1).

#### Ground truth maps

We used the National Land Cover Database (NLCD) to associate Landsat pixels with land cover classes (Yang et al., 2018). The NLCD for our study region included 15 land cover classes (Supplementary Materials 1.2). The NLCD updates land cover classifications approximately every three years. We obtained historical NLCD maps from seven years that fell within the 21-year span of our study: 2001, 2004, 2008, 2011, 2013, 2016, and 2019. We then added the 3,345 leafy spurge occurrence points by reclassifying existing NLCD maps based on reported survey dates (e.g. if an occurrence was reported in 2018, 2019, or 2020, we updated the 2019 NLCD map; Supplementary Methods 1.2; Table S1).

#### Temporal convolutional neural network (TempCNN) architecture and training data

We developed a temporal convolutional neural network (TempCNN) (Pelletier et al., 2019) using Landsat spectral data from the 21 years (Figure 1). A TempCNN is a supervised deep learning algorithm that can learn from spectral and phenological information in a satellite image time series (Allred et al., 2021; Pelletier et al., 2019). To train the TempCNN, we constructed a dataset that included all community science leafy spurge occurrences and randomly sampled points from across Minnesota. We divided the timeframe (2000-2020) into seven intervals of three years (e.g. 2000-2002, 2003-2005, etc.). For each interval, we included leafy spurge occurrences recorded during that timeframe and one hundred thousand randomly sampled pixels from the study region (Tables S1; S2). For each sampled pixel, we constructed a data vector that included: latitude, longitude, land cover class, and spectral data from nine seasonal windows (i.e., March/April, May/June, and July/August for the three consecutive years of the temporal interval). Land cover class was obtained from the NLCD map that was updated in one of the three years included in an interval. We repeated this process over the seven intervals from 2000-2020, yielding approximately seven hundred thousand samples for model training (for additional details see Supplementary Materials 1.3). This data structure allowed the neural network to learn from spectral, phenological, and spatial features.

**Figure 1.**
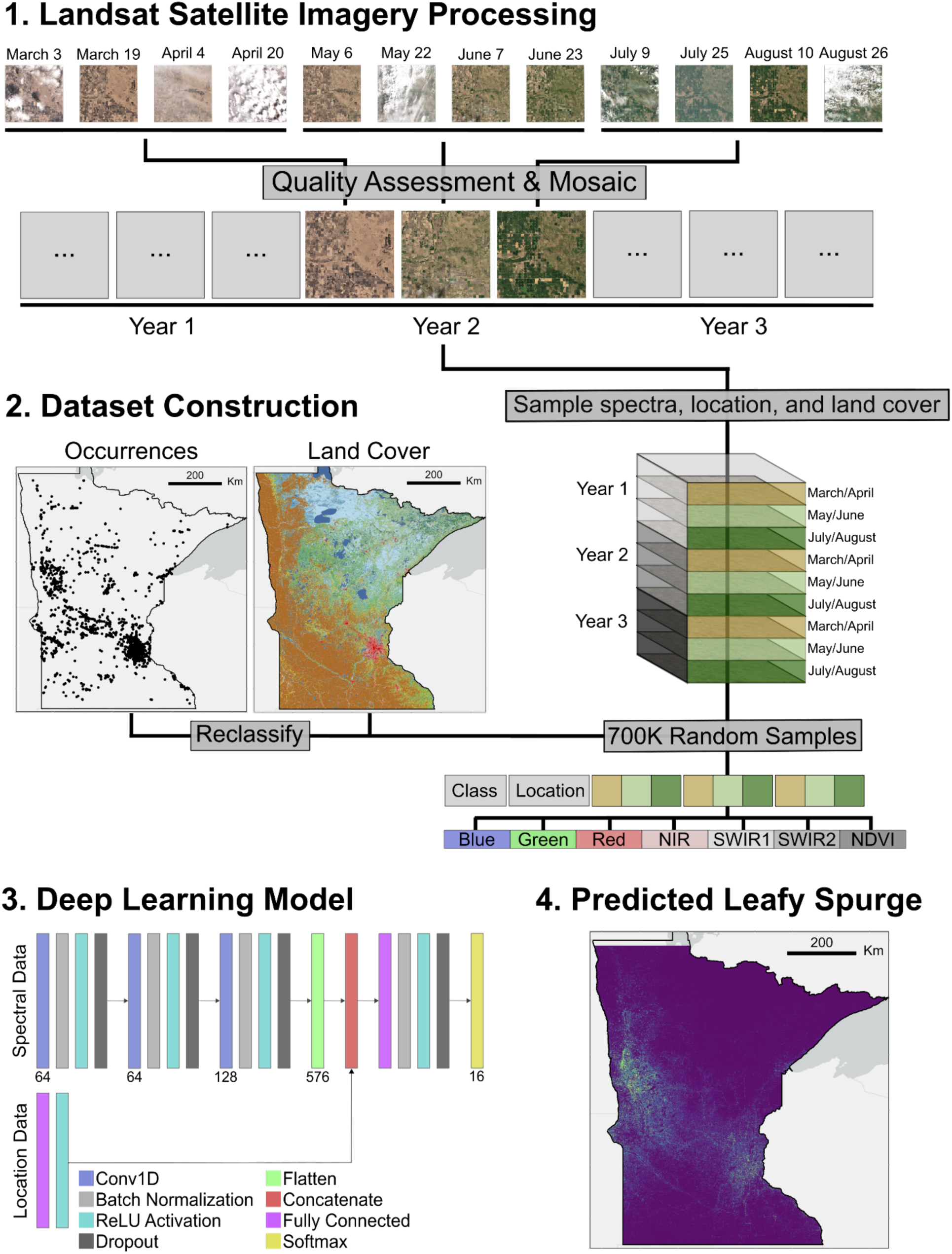
Overview of remote sensing models. 1) Landsat satellite image time-series mosaics were processed with Google Earth Engine. Landsat imagery (2000-2020) included three two-month periods per year: March-April, May-June, and July-August. 2) We constructed a data vector for each occurrence record used to train the temporal convolutional neural network (TempCNN). Community science occurrences for leafy spurge were obtained from publicly available databases. In addition, approximately 700,000 random points were sampled from the National Land Cover Database between 2000 and 2020. The data vector included the land cover classification from the National Land Cover Database, location (lat/lon), and spectral information from a three-year time frame centered on a temporal interval of interest. 3) A temporal convolutional neural network (TempCNN) was used to learn the relationship between land cover class and the three years of spectral data and location information. 4) After training and evaluating the TempCNN model, leafy spurge and land cover classes were predicted across the state of Minnesota.

**Figure 2.**
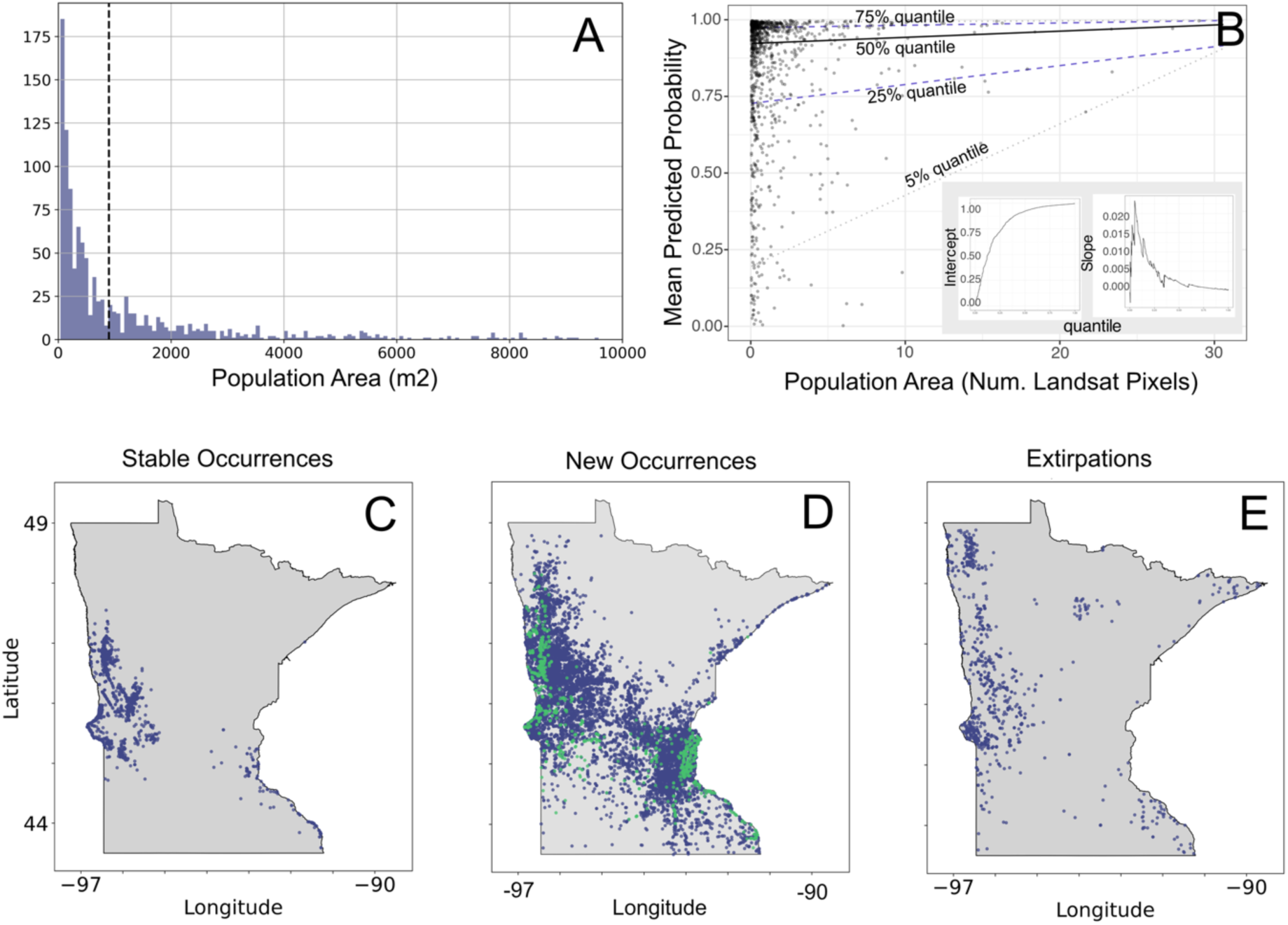
(A) Population area distribution of leafy spurge validation occurrences collected 2021-2023. (B) Relationship between predicted probability of leafy spurge from the TempCNN model and population area, presented as number of pixels (one pixel = 900m^2^). The lines represent quantile regression relationships. The inset shows the relationship between the quantile modeled using quantile regression and the intercept (left) and slope (right) of those relationships. (C-E) TempCNN predictions of temporal change in leafy spurge distribution. (C) Stable leafy spurge occurrences from 2000-2020; predicted with ≥0.5 probability at all time periods (N=3,358 points). (D) New leafy spurge occurrences. Green points appeared by 2006-2008 (N=1,286 points). Blue points appeared after 2014 (N=17,609 points). (E) Extirpated leafy spurge populations were present in 2000-2005 but absent after (N=4,918 points).

Our model architecture included a combination of 1D convolutional layers, dropout layers, and fully connected layers. Convolutional layers were used to learn the spectral and phenological profiles of each land cover class, dropout layers minimized model overfitting, and fully connected layers passed information between layers. In the final step, the model outputs pixel-specific predictions of land cover class probabilities using a softmax layer (i.e. the response for every pixel is a vector of 16 numbers corresponding to the probability of each land cover class, range: 0-1). Finally, to generate pixel classifications to test model accuracy, we used an argmax layer that selected the land cover class with the highest probability as the class assignment (for additional details see Supplementary Materials 1.4).

#### TempCNN training, evaluation, and prediction

We split our dataset into training, validation, and testing sets using an 80/10/10 ratio (Table S2). We trained the model for 100 epochs using the Adam optimizer and categorical cross-entropy loss function (Figure S1). We applied class weights during training to assist with classifying imbalanced data (Table S2; for additional details see Supplementary Materials 1.5). We then evaluated model performance on the testing set by calculating accuracy, specificity (i.e. true negative rate), sensitivity (i.e. true positive rate), and the F2 score [a measure of performance that accounts for model precision (proportion of predicted positives that are correct) and sensitivity] for each class (for additional details see Supplementary Materials 1.6).

We predicted leafy spurge at each of our seven temporal intervals at a 30-meter resolution across Minnesota, totaling approximately 250 million pixels (Figure 1, for additional details see Supplementary Materials 1.7).

In addition to traditional confusion matrix-based evaluations of our model, we also wanted to gauge the predictive capacity and underlying probability distributions of the softmax layers produced by our TempCNN. To do this, we compared the model softmax outputs from our most recent prediction (2018-2020) for each land cover class to (1) an independent occurrence dataset for leafy spurge (2021-2023) not used for model training and (2) the NLCD 2021 dataset (i.e. the 15 land cover classes, not including leafy spurge). The leafy spurge dataset consisted of recent leafy spurge occurrences from Minnesota (2021-2023; N = 1,018; Table S1). For a subset of the records (N = 940), population area (measured by the area of a spatial polygon and converted to m^2^) was estimated by management professionals (Figure S2). We generated a leafy spurge prediction across Minnesota using Landsat imagery from 2018-2020 and determined if the recently reported occurrences (2021-2023) were located in pixels with a high predicted leafy spurge probability. We also examined whether the predicted probability of leafy spurge was associated with population area (N = 940) using quantile regression (for additional details see Supplementary Methods 1.7). For other land cover classes, we used the NLCD 2021 dataset because it was not used in model training. We randomly sampled 10,000 points per land cover class from the NLCD 2021 dataset and extracted the predicted probability values for these points from the associated softmax land class prediction. We visualized the histogram of probability values and estimated the median predicted probability for each land cover class (for additional details see Supplementary Methods 1.7).

#### Thresholding predictions for total area invaded

We transformed TempCNN softmax layer predictions of leafy spurge into discrete occurrence points. We chose a probability threshold for a positive leafy spurge presence of 0.5 because it indicated when the probability of leafy spurge was greater than the sum of the probability of all other classes. It may be taken as a conservative threshold estimate because it was greater than the threshold required to obtain 75% sensitivity of occurrences reported by community scientists (threshold = 0.38 estimated at the 2009-2011 midpoint prediction period; Figure S3). We applied this threshold to all timeframes to determine the geography and timing of leafy spurge occurrences and used the 2009-2011 TempCNN prediction map to generate a remotely-sensed occurrence dataset for spatial bias analyses.

#### Extent of invasion and climate trends

We tested the hypothesis that remotely-sensed estimates of the area invaded were affected by climate variation. Specifically, if drought reduces plant biomass and flower production, the remote sensing model may predict that fewer pixels were invaded. We used the Palmer Drought Severity Index (PDSI) (Palmer, 1965) to capture variation in temperature, precipitation, and soil moisture, which influence the timing and magnitude of leafy spurge flowering (Selleck et al., 1962). PDSI values range from -15 to +15, with lower values indicating more severe drought. To align with our remote sensing predictions, we averaged PDSI values over three-year intervals (e.g., 2000-2002) and partitioned the data into nine geographic regions based on the National Oceanic and Atmospheric Administration’s climate divisions. We quantified invaded area for each region and timeframe by applying the 0.5 presence threshold to the corresponding TempCNN leafy spurge softmax prediction.

We used a generalized linear model to test whether predicted invasion extent was related to drought severity and geographic region. We quantified leafy spurge extent as the number of positive leafy spurge pixels in each of the nine regions (Figure 3). We used the ‘glm’ function and specified family as ‘quasipoisson’ due to overdispersion in the data. Drought severity (PDSI), geographic region, and their interaction were independent predictors. We assessed significance of model terms using analysis of deviance with X^2^ tests from the ‘Anova’ function from the ‘car’ package and quasi-F tests from the base ‘anova’ function in R (Dunn and Smith, 2018). We estimated mean values using the ‘emmeans’ R package (Lenth 2025).

**Figure 3.**
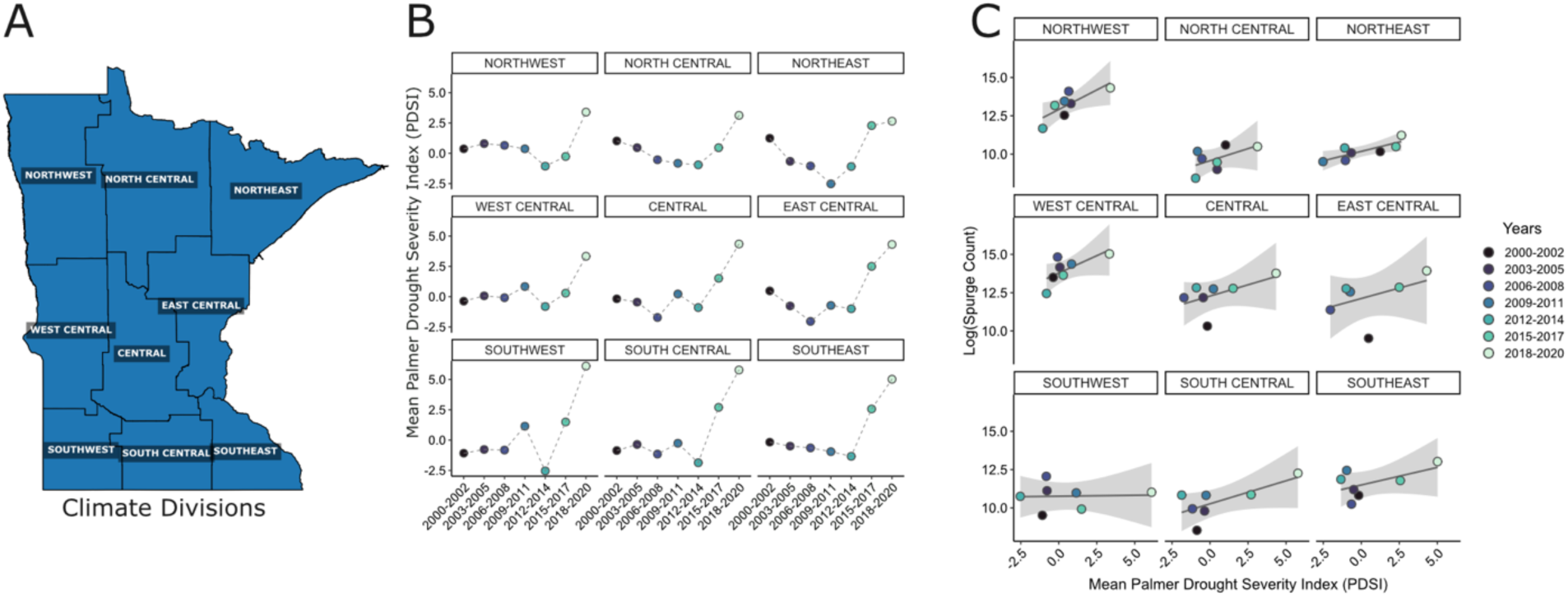
Regional climate variation shapes the predicted extent of leafy spurge invasion across Minnesota. (A) Map of Minnesota’s nine climate divisions, defined by the National Oceanic and Atmospheric Administration’s National Centers for Environmental Information. (B) Mean Palmer Drought Severity Index (PDSI) values averaged over three-year periods from 2000 to 2020 for each climate division. Negative values indicate drought conditions, while positive values reflect wetter periods. (C) Relationship between leafy spurge invasion extent and mean PDSI, with data faceted by climate division. The y-axis shows the log-transformed invasion extent predicted by the TempCNN model; the x-axis shows mean PDSI values across corresponding three-year periods.

### Geographic patterns, spatial bias, and timing of invasion

#### Geography and timing of new occurrences and extirpations

We examined the geography and timing of new occurrences, extirpations, and continually present populations from 2000 to 2020. We categorized new occurrences into two time periods: early occurrences and late occurrences. We defined early occurrences as pixels where leafy spurge was absent in the first two prediction intervals (i.e., 2000-2002 and 2003-2005) but present in all subsequent five periods (i.e., 2006-2006 through 2018-2020). Similarly, we defined late occurrences where leafy spurge was absent in the first five prediction intervals and present in the last two periods. Extirpations included pixels where leafy spurge was present in the first two predictions and absent in all five subsequent periods. Continually present populations were those pixels where leafy spurge remained present across all seven predictions from 2000 to 2020. For all categorizations, we used the 0.5 threshold for positive presences and a conservative threshold for absence (0.15).

#### Estimating population size changes through time

We estimated population change from 2000 to 2020 for individual pixels using linear regression. For each Landsat pixel across the study region, we regressed the probability of leafy spurge for seven intervals from 2000-2020 on year. The slope of the regression for each pixel served as a proxy for the rate of population change. Positive slopes (> +0.5 SD from mean) were taken to indicate growth, negative slopes (< -0.5 SD from mean) were taken to indicate decline, and slopes near zero indicated stability or fluctuations through time.

#### Spatial overlap and autocorrelation of occurrence datasets

We measured spatial autocorrelation for the community science and remotely-sensed occurrence datasets independently (N = 1,771; remotely-sensed occurrence dataset description below, see *Occurrence weighting for SDMs*; Figure 5). We used Moran’s I to determine if occurrences were more clustered than random expectation. We calculated Moran’s I using the ‘moran.mc’ function in the ‘spdep’ R package v1.3-4 (Bivand et al., 2015). We determined if the value was significantly greater than random at a significance level of 0.05 using permutation tests (n = 999).

**Figure 4.**
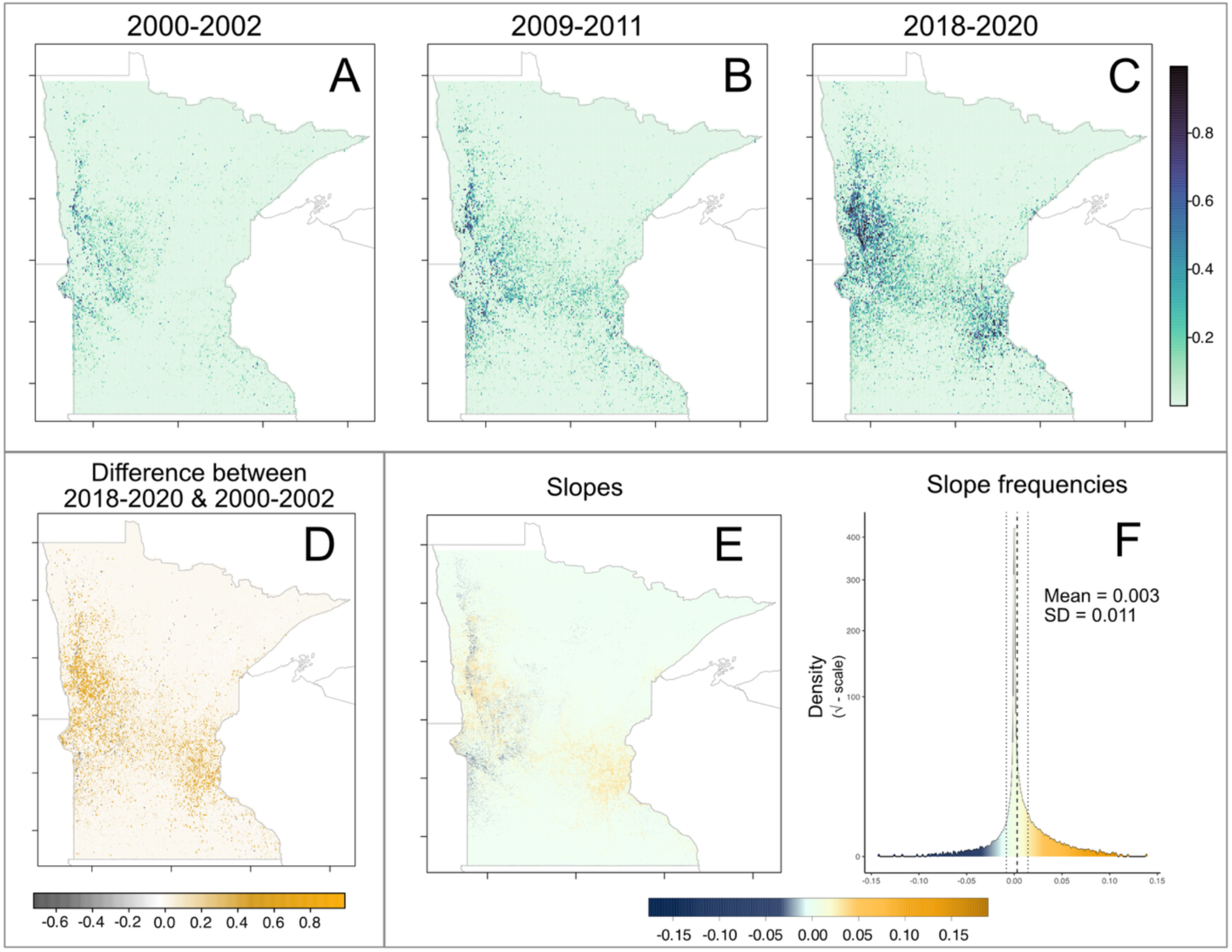
Predicted probability and population dynamics of leafy spurge for individual pixels from the TempCNN. (A-C). Probability of leafy spurge for (A) 2000-2002, (B) 2009-2011, and (C) 2018-2020. (D) The difference between the 2018-2020 and 2000-2002 prediction. Orange pixels indicate positive values and grey pixels indicate negative values. (E) Slope values from regressions of predicted probability versus time period. (F) Histogram of the slope values. The dashed line indicates the mean slope and the dotted lines are ± standard deviation.

**Figure 5.**
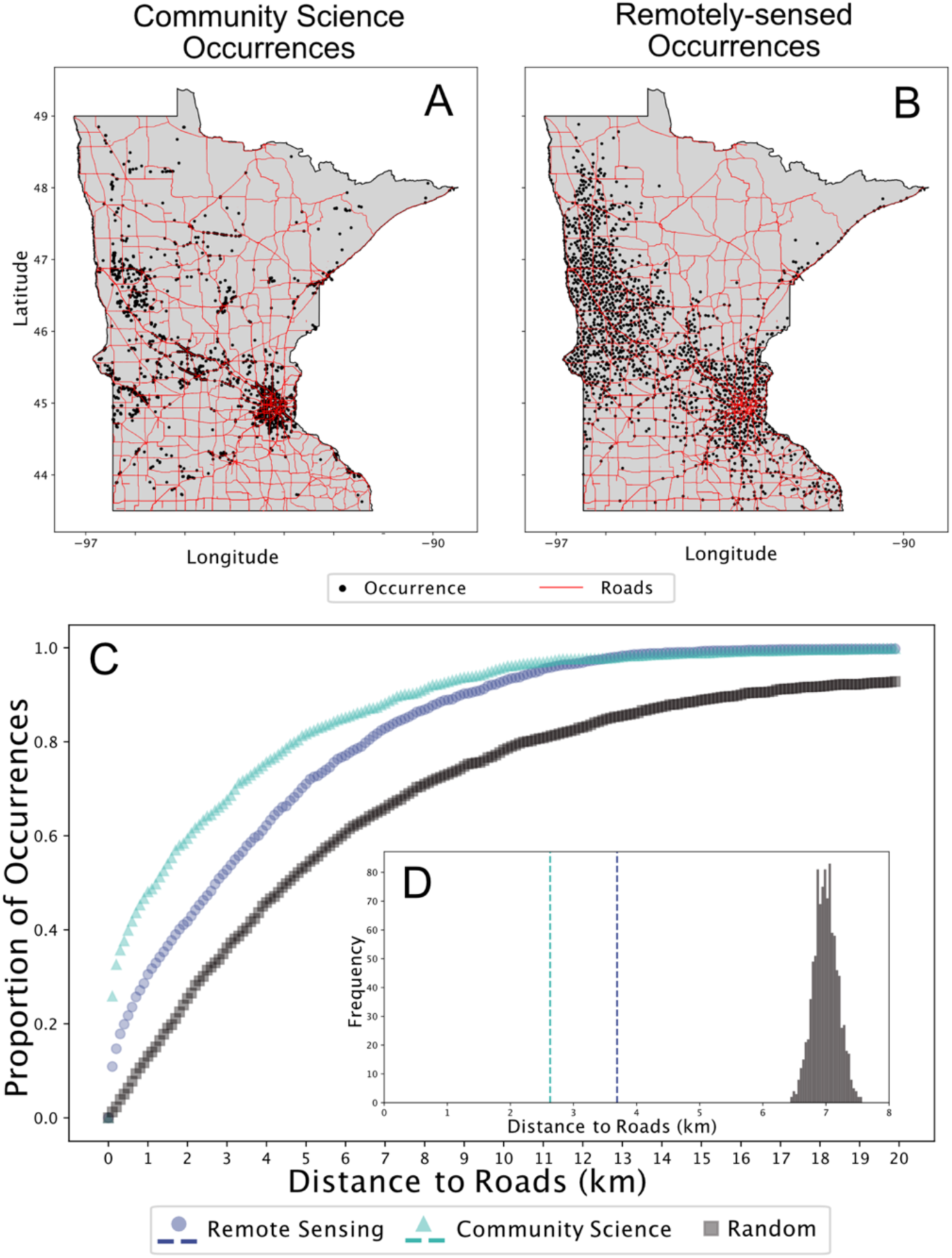
Leafy spurge occurrences used for testing spatial sampling bias and SDM development. (A-B) Mapped occurrences (black dots) and major roads (interstates and highways; red lines). (A) Community science (CS) occurrences from 2000-2020 (N = 1771). (B) Random sample of remotely-sensed (RS) occurrences (N = 1771) from the TempCNN. (C). Relationship between the cumulative proportion of occurrence records and distance to the nearest road. (D-inset) Comparison of the mean distance to a roadway for CS occurrences (2.63 km), RS occurrences (3.69 km), and a distribution of randomly drawn occurrences (N=999).

To quantify the extent to which the community science and remotely-sensed occurrences co-localized, we quantified spatial overlap between occurrences from the two datasets using the Jaccard index [range: 0 (no overlap) to 1 (complete overlap)]. We used a 0.1-degree latitude/longitude grid to create binary presence-absence maps for both datasets. We then calculated the Jaccard index as the intersection over union between both raster datasets using the ’raster’ R package (Hijmans et al., 2015).

#### Testing for roadside bias

Roadside surveys are a major contributor to sampling bias in occurrence datasets (Geurts et al., 2023). To estimate the potential for roadside bias in our datasets, we measured the distance to primary and secondary roads for both community science and remotely-sensed occurrences (as above N = 1,771; Figure 5) using the U.S. Census Bureau’s MAF/TIGER Database (for additional details see Supplementary Materials 1.8). We used a k-d tree nearest neighbor search to calculate the shortest distance between occurrence points and road segments. For comparison, we generated a null expectation by calculating distances to the nearest road for an equal number of randomly selected points (N = 999 permutations) within Minnesota. We compared the mean distance to roads for both occurrence sets and determined if either value was less than random expectation at the 0.05 level of significance (one-tailed test).

### Species distribution modeling

#### Modelling overview

We developed a set of species distribution models (SDMs) to examine whether the use of remotely-sensed occurrences and demographic weighting can improve predictions of climate suitability. We evaluated three weighting schemes for occurrences: unweighted (traditional), weighted by the model’s confidence in the prediction from the remote sensing model (e.g. high confidence could be caused by high density), and weighted by slope of population size change through time. Because we were interested in comparing how the occurrence datasets perform across modeling platforms, we built SDMs using two commonly-used approaches, Maxent and Boosted Regression Trees (BRT) (Elith et al., 2008; Kass et al., 2021; Phillips et al., 2017), in R v.4.3.1 (R Core Team, 2023). Spatial data processing was accomplished using the ’rgdal’ (R. Bivand, Keitt, et al., 2015), ‘rgeos’ (M. R. Bivand, 2018), ’sp’ (Pebesma & Bivand, 2013), and ’raster’ packages (Hijmans et al. 2015).

#### Environmental data

We downloaded three bioclimatic variables at a 30 arcsecond resolution (∼1km) from WorldClim v.2.1 (worldclim.org): mean temperature of the warmest quarter (Bio 10), mean temperature of the coldest quarter (Bio 11), and annual precipitation (Bio 12) (Figure S4). These axes of climate variation are relevant to key life history stages of leafy spurge in Minnesota (Briscoe Runquist & Moeller, 2024; Lake et al., 2023). Warm season temperatures affect seed germination, growth, and phenology (Wolkovich et al., 2013), cold season temperatures influence winter survival and damage (Chapman et al., 2017), and precipitation influences growth and reproduction (Petitpierre et al., 2017).

#### Occurrence weighting for SDMs

We fit SDMs using both community science and remotely-sensed occurrences. For SDM development, the starting remotely-sensed occurrence dataset combined the continuous and new early and late occurrence points (see *Geography and timing of new occurrences and extirpations*). The community science and remotely-sensed datasets were thinned using grid sampling to match the raster resolution of the environmental data. After thinning, 1,771 and 22,204 occurrence records remained, respectively. We randomly sampled 1,771 occurrences from the thinned remotely-sensed dataset to generate the final remotely-sensed occurrence dataset (Figure 5).

We developed unweighted (traditional) and weighted SDMs to evaluate the extent to which remotely-sensed demographic information improved the accuracy of SDMs. We weighted occurrences by: (1) confidence in the leafy spurge prediction from the TempCNN and (2) population size change over the 21-year study (the slope from linear regressions of the probability for leafy spurge on time).

#### Modeling algorithms

For Maxent models, we used the ENMeval package to generate a collection of nine models that reflected a set of commonly used hyperparameter combinations. We did this separately for each occurrence dataset and weighting combination, resulting in six groups of models. Assessing multiple models in each group also allowed us to evaluate model stability, model discrimination, and the potential for overfit (Radosavljevic & Anderson, 2014). ENMeval allows the user to vary the feature classes used (linear (L), quadratic (Q), polynomial (P), threshold (T) and hinge (H)) and the smoothing regularization parameter (beta). We developed models using three feature class combinations that vary in complexity (LQH, LQHP, LQHPT) and three beta parameters that vary regularization (1, 2, 3).

To develop the Maxent models, we generated 10,000 random background points to characterize the distribution of environmental conditions across Minnesota. We used four-fold cross-validation by randomly partitioning occurrences and background points into spatial blocks using ‘checkerboard 2’ partitioning to reduce spatial autocorrelation among cross-validation blocks (Figure S5). We then ran the ENMeval function using the ‘maxnet’ implementation of the maximum entropy model.

To weight occurrences in Maxent models, we generated “pseudopresences” for models using probability quantiles from data output from our remote sensing model. The “pseudopresences” were records added to the dataset that replicated the environmental values of occurrences where the number of replicates of an occurrence indicated the weighting (Eckhart et al., 2011). For weighting by the remote sensing model’s probability estimates, occurrence records with a probability below the 25th percentile were represented once in the dataset, records between the 25th and 50th were represented twice, records between 50th and 75th were represented three times, records between the 75th and 99th were represented four times, and those above the 99th percentile were represented five times. To generate weighting for demographic growth estimates, we repeated this process but used slopes to generate the quantile distribution.

For BRT models, we also generated a collection of nine models for each occurrence dataset and weighting combination. This allowed us to represent a set of commonly used hyperparameters and to assess model stability, discrimination and overfit. We selected BRT hyperparameters for each set of models based on practices outlined in Elith et al. (Elith et al., 2008). We varied tree complexity (2, 3, or 5) and learning rate (three values between 0.001 and 0.01). The algorithm required that weighting values be positive, so in population size change models, we assigned randomly generated very small positive values (mean: 1e-4) for records with negative slopes (CS: N = 416; RS: N = 74). During model training we set the bag fraction to 0.8. Learning rates varied slightly to ensure that we produced well-regularized models that reach an asymptote in model deviance within 10,000 trees.

To develop the BRT models, we generated absence points equal to the number of presences. For community science models, we sampled pseudoabsences from across the sample area as is common practice in SDM studies (Elith et al., 2010). For the remotely-sensed dataset, we sampled absence points from regions where the remote sensing models predicted leafy spurge at less than five percent probability (Figure S6). We built models using the ‘gbm.step’ function from the ’dismo’ and ‘gbm’ packages to select the model with the optimized deviance for the hyperparameters chosen (Greenwell et al., 2019; Hijmans et al., 2017).

#### Model evaluation metrics

We evaluated SDM performance using multiple metrics. We used the average Area Under the Curve (AUC) across four cross-validation folds to assess model discrimination (Kass et al., 2021; Radosavljevic & Anderson, 2014). AUC measures how well the model distinguishes between presence and absence, with values ranging from 0.5 (random) to 1.0 (excellent). We report AUC at the threshold where the true positive rate minus the false positive rate was minimized. To detect model overfitting, we calculated the average difference between the calibration and evaluation AUC values across the four cross-validation folds. The AUC difference quantifies the degree to which the model’s predictive performance on the training data (calibration) exceeds its performance on unseen data (evaluation), where higher values indicate a greater extent of overfitting (Radosavljevic and Anderson 2014).

AUC can be misleading due to its equal weighting of omission and commission errors, which may be especially problematic for invasive species where omission errors may be more costly (Leroy et al., 2018; Lobo et al., 2008). Therefore, we also calculated sensitivity, which measures the proportion of correctly predicted occurrences. We calculated sensitivity using the 1,018 occurrence records collected from iNaturalist and EddMaps between 2021-2023 (Figure S2). These records were not used during model training and represent an independent dataset composed of the type of data (e.g. new occurrence records) that SDMs are often developed to predict. Sensitivity was evaluated at a threshold that maximized the difference between the true positive rate and the true negative rate. One of the potential strengths of remote sensing is in detecting occurrences in difficult-to-survey, remote regions. To evaluate model sensitivity in rural compared to urban areas, we divided our data between the Twin Cities Metropolitan region and outlying rural areas (Figure S7).

We tested for significant differences in AUC, overfitting (AUC difference), and sensitivity among different data sources (community science vs. remote sensing) and weighting schemes (traditional, probability, slope) separately for model types (Maxent vs. BRT). For each evaluation metric, we used ANOVA with occurrence data source, weighting, and their interaction as independent variables. We checked all models to ensure that they conformed to assumptions of normality and homogeneity of variance in residuals. We assessed significance using the ‘Anova’ function from the ‘car’ package in R using type II sums of squares (Fox & Weisberg, 2018). We performed Tukey’s tests to evaluate the significance of pairwise comparisons using the ‘emmeans’ R package (Lenth, 2025).

## RESULTS

### Remote sensing accurately classified leafy spurge and land cover classes

The TempCNN predicted leafy spurge with high accuracy (97%), comparable to the model’s predictive accuracy for other classes (Table S3). When evaluated using testing data, leafy spurge was predicted with high specificity (97%) and moderate sensitivity (65%). The leafy spurge F2 score was 32%, similar to F2 scores of other classes (Table S3). The moderately low F2 scores may be indicative of low precision from false positive identifications.

Misclassifications among leafy spurge, grasslands/herbaceous, pasture/hay, and low-intensity developed land cover classes were among the most common sources of error (Figure S8).

We found agreement between our predicted land cover classes from 2018-2020 and the 2021 NLCD dataset, indicating that the TempCNN generalized beyond the training data (Figure S9). Recent leafy spurge occurrences (2021-2023) were also well captured by the TempCNN. From the softmax probability layer predictions, the median probability of leafy spurge occurrences was 48%, which was similar to the median predicted probabilities of other classes in the TempCNN model – e.g. deciduous forests (46%) and developed high-intensity (54%) land cover classes (Figure S9).

The mean and median probability of detection were high across population size variation in our independent validation dataset (all means > 0.80; all medians > 0.92; Figure 2). However, the quantile regression showed that small populations had higher variance in detection probability than larger populations. Similarly, the median quantile had a high intercept (0.92) and shallow slope (0.002) whereas the 5% and 25% quantiles had lower intercepts (0.19 and 0.73) and steeper slopes (0.23 and 0.006, respectively; Figure 2).

### Temporal dynamics of invasion and the effects of drought on remote sensing predictions

The invaded area expanded from 1,067 km² in 2000–2002 to 7,156 km² in 2018–2020 (Table S4). Invasion extent also fluctuated over time, with declines in 2009–2011 (3,124 km^2^) and 2012–2014 (1,372 km^2^) before increasing in subsequent years (Table S4). The extent of invasion was significantly influenced by drought severity and region, which explained 10.2% and 84.7% of the model deviance, respectively (Table S5). Greater drought severity was associated with reduced detection by the TempCNN (and lower invasion extent; (Figure 3). The relationship was strongest in west-central and northwest Minnesota (Figure 3). These findings suggest that climate variation influences the performance of leafy spurge, which in turn affects the capacity to detect it from satellite images.

### Geography and timing of new occurrences and extirpations

Leafy spurge was already well established in Minnesota prior to the timeframe of our study. Populations were most abundant in the prairies and grasslands of west-central Minnesota, with smaller but notable presence in urban areas (Figures 2 & 4). In the timeframe of our study, we observed new areas of invasion across most of the state except the far northern and southern regions (Figures 2 & 4). Between 2000-2010, newly-detected populations were primarily concentrated in prairies/grasslands near urban areas on the western (Fargo-Moorhead) and eastern sides of the state (Minneapolis-St. Paul). Between 2010-2020, substantially more spatial spread radiated from urban to rural areas, including many new occurrences in central Minnesota. Extirpations were primarily detected in western prairies/grasslands but were absent from urban areas (Figures 2 & 4).

### Population dynamics of leafy spurge through time

For individual pixels, the mean slope of the relationship between leafy spurge probability and time was 0.003 (std dev: 0.001), indicating an overall increase in leafy spurge across Minnesota (Figure 4; Table S4). We detected population growth in 13.1% of the state (> +0.5 SD from mean). Population growth was observed primarily in urban areas (i.e., Minneapolis-St. Paul and Fargo-Moorhead) and prairie or grasslands (i.e., west-central Minnesota) (Figure 4). We detected population decline in 8.4% of the state (< -0.5 SD from mean). Population decline was observed primarily in the boreal region (i.e., northeastern Minnesota) and portions of western Minnesota (Figure 4).

### Spatial biases in community science occurrences

The spatial overlap between remotely-sensed and community science occurrences was modest (Jaccard’s Index = 0.36) (Figure 5). Spatial autocorrelation was significant and similar for the community science (Moran’s I = 0.221; *p < 0.001*) and remotely-sensed occurrences (Moran’s I = 0.242; *p < 0.001*). These results indicate that occurrences tend to be locally clustered in both datasets.

Remotely-sensed occurrences were more likely to be found near roads than random expectation (Figure 5; *p* < 0.001). Remotely-sensed occurrences were an average of 3.7 km from roads and 50% of occurrences were within 3 km of a road (Figure 5). Community science occurrences were even more likely to be associated with roads than remotely-sensed occurrences (Figure 5). On average, community records were 2.6 km from roads and 50% of occurrences were within 1 km of a road. This result indicates spatial bias in the community science dataset that is likely due to sampling.

### Remotely-sensed occurrences improved SDM accuracy especially in non-urban regions

The occurrence dataset (community science vs. remotely-sensed) had the largest influence on SDM evaluation metrics (Figure 6; Table 1) and climate suitability predictions (Figures 7, S10). Models fit using remotely-sensed occurrences had higher discrimination scores in Maxent and BRT models (Reported values are mean ± SE) (AUC: Maxent: CS: 0.766 ± 0.001; RS: 0.772 ± 0.001; BRT: CS: 0.864 ± 0.002; RS: 0.946 ± 0.002). In addition, SDMs built with community science data were always more overfit relative to SDMs built with remotely-sensed occurrences (AUC difference; Maxent: CS: 0.055 ± 0.001; RS: 0.015 ± 0.001; BRT: CS: 0.059 ± 0.001; RS: 0.017 ± 0.001).

**Figure 6.**
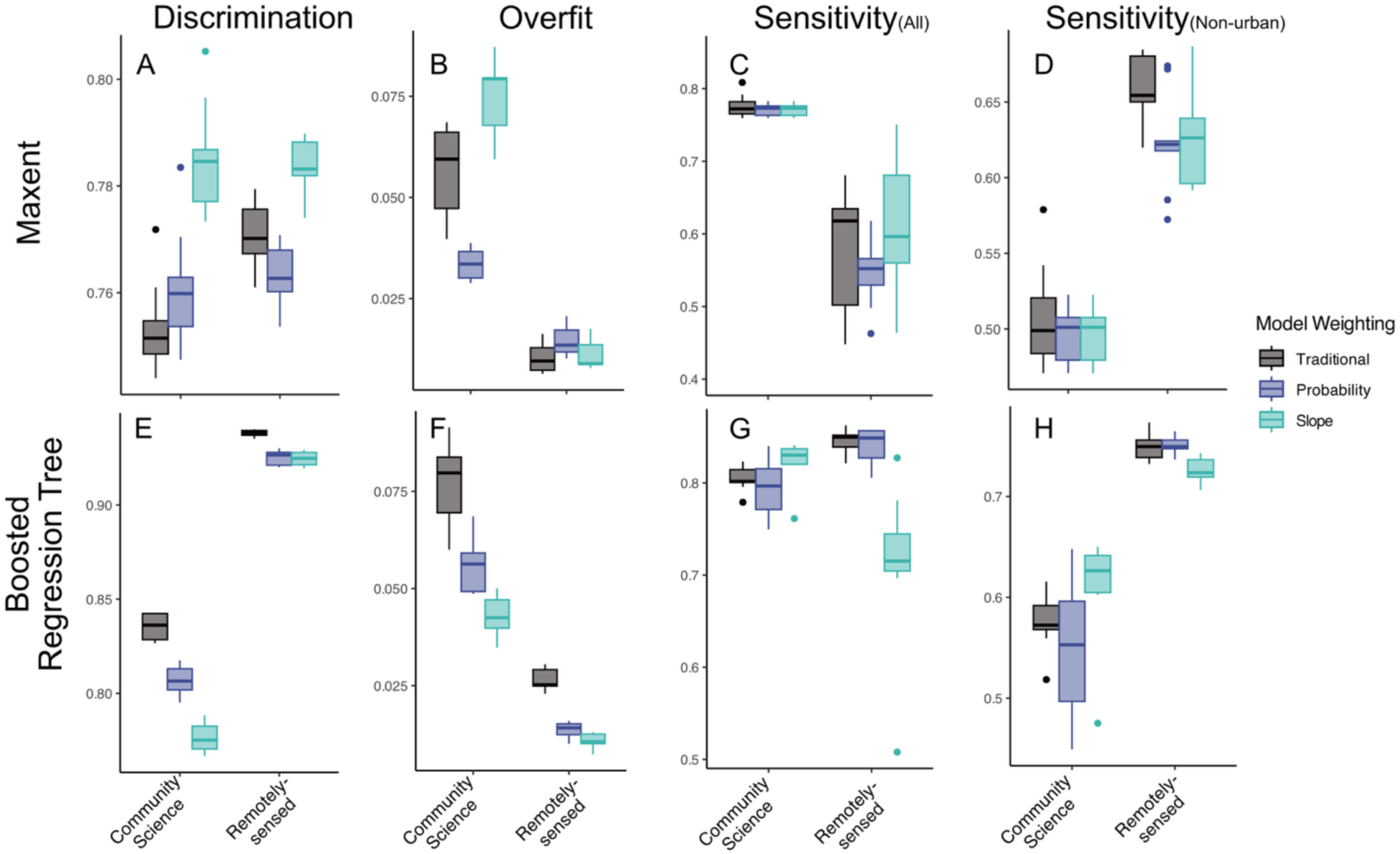
Comparison of species distribution model evaluation metrics. Maxent and Boosted Regression Tree models were developed using either community science (CS) or remotely-sensed (RS) occurrences. Occurrences were also either unweighted (traditional) or weighted by predicted probability or population growth/decline. (A & E) Model discrimination is the average AUC across four cross-validation folds, where higher values indicate higher discrimination ability. (B & F) Model overfit measures the average difference between the calibration and evaluation AUC values, with higher values indicating greater overfit. (C & G) Sensitivity to all independent leafy spurge records sampled from 2021 – 2023, where higher values indicate higher performance. (D & H) Sensitivity to independent records sampled from 2021 – 2023 from non-urban areas, where higher values indicate higher performance.

**Figure 7.**
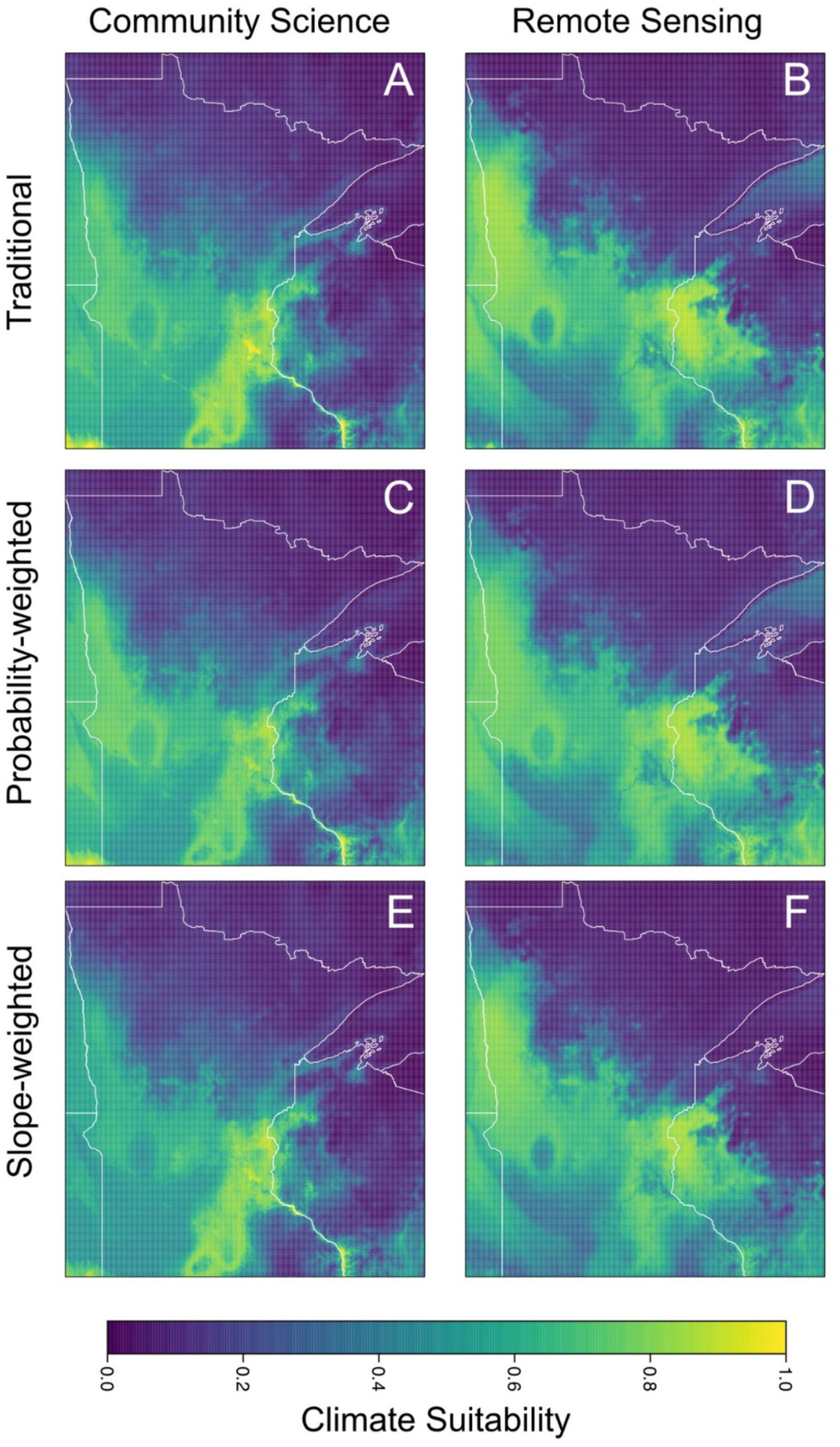
Maxent model projections of climate suitability from models built using either community science or remotely-sensed occurrences and weighting of occurrences (traditional, probability, and slope = population growth/decline). Each model projection is the average of nine models built with different combinations of hyperparameters.

**Table 1.** ANOVA tables to test for difference in four SDM evaluation metrics (AUC, Overfit, Sensitivity – All, and Sensitivity – Non-urban) due to using different data sources (remotely-sensed or community science) and/or weighting schemes (traditions, probability, slope) to build SDMs. The table presents results for two algorithms: Maxent (left) and Boosted Regression Trees (BRT; right). Significant P-values (*P* <0.05) shown in bold.

|  | Maxent |  |  | BRT |  |  |
| --- | --- | --- | --- | --- | --- | --- |
|  | df | F | P | df | F | P |
|  | AUC |  |  |  |  |  |
| Dataset | 1 | 7.4 | **0.009** | 1 | 749.2 | **<0.001** |
| Weighting | 2 | 41.0 | **<0.001** | 2 | 140.8 | **<0.001** |
| Dataset*Weighting | 2 | 6.0 | **0.005** | 2 | 37.8 | **<0.001** |
|  | Overfit |  |  |  |  |  |
| Dataset | 1 | 582.1 | **<0.001** | 1 | 658.3 | **<0.001** |
| Weighting | 2 | 38.3 | **<0.001** | 2 | 81.3 | **<0.001** |
| Dataset*Weighting | 2 | 55.8 | **<0.001** | 2 | 10.1 | **<0.001** |
|  | Sensitivity - All |  |  |  |  |  |
| Dataset | 1 | 164.8 | **<0.001** | 1 | 0.5 | 0.482 |
| Weighting | 2 | 1.3 | 0.291 | 2 | 10.6 | **<0.001** |
| Dataset*Weighting | 2 | 1.2 | 0.297 | 2 | 21.9 | **<0.001** |
|  | Sensitivity - Non-urban |  |  |  |  |  |
| Dataset | 1 | 301.0 | **<0.001** | 1 | 255.8 | **<0.001** |
| Weighting | 2 | 3.0 | 0.057 | 2 | 1.3 | 0.2782 |
| Dataset*Weighting | 2 | 0.72 | 0.494 | 2 | 7.0 | **<0.001** |

When evaluating SDMs using an independent occurrence dataset, SDM sensitivity was dependent on geography (Figure 6). Across Minnesota, including the highly invaded urban Twin Cities region, SDMs built using community science occurrences generally had higher sensitivity scores for both modeling algorithms, although it was only significant for Maxent models (Maxent: CS: 0.79 ± 0.01; RS: 0.56 ± 0.01; BRT: CS: 0.81 ± 0.01; RS: 0.80 ± 0.01). Conversely, in non-urban areas, SDMs built using remotely-sensed occurrences had higher sensitivity scores (Maxent: CS: 0.50 ± 0.01; RS: 0.69 ± 0.01; BRT: CS: 0.58 ± 0.01; RS: 0.74 ± 0.01).

SDM occurrence weighting schemes (traditional, probability-weighted, slope-weighted) had more subtle effects on evaluation metrics and the effects were more pronounced for BRT models (Figure 6; Table 1). When significant, weighting affected the relative order and variance among model metrics within the two different dataset types. Although weighting did not always improve model metrics when compared to traditional SDMs, SDMs were generally less overfit with similar sensitivity values when using remotely-sensed data (Figure 6).

## DISCUSSION

Accurate forecasts of geographic range shifts with climate change are important for conservation of rare species and management of invasive species. Predictive distribution models rely on unbiased occurrence datasets to determine occurrence-environment relationships but publicly-available datasets nearly always suffer from spatial bias and lack of information on demography. In this study, we used a 21-year time series of Landsat imagery coupled with deep learning to detect leafy spurge and assess spatial and population changes over 225 thousand square kilometers in Minnesota, USA. Our analyses show that the area invaded by leafy spurge increased by as much as 570% over 21 years. Invasion during this period involved range expansion and range infilling in two distinct areas in Minnesota that differ in climate. Remotely-sensed occurrences were biased to roadsides, albeit less biased than community science occurrences. These results suggests that factors other than climate (e.g. disturbance) may be particularly important in driving invasion. Including remotely-sensed occurrences and population growth in SDMs reduced overfitting and improved predictions compared to models using community data alone. The greatest improvement in model fit occurred at the western range edge and near urban centers, where growth rates were highest. Our findings indicate that remote sensing can dynamically monitor invasion, reduce spatial bias in occurrences, and improve species distribution models.

### Factors modulating detection and inferences about invasion

Leafy spurge was detectable from satellite imagery due to its distinct phenology (emergence, flowering, and senescence) and spectral signal (yellow-green flowering bracts). While previous remote sensing surveys were limited to local and regional scales (10 to <1000 square kilometers) (Hunt & Parker Williams, 2006; Lake et al., 2022; Mattilio et al., 2023; Mitchell & Glenn, 2009; Mladinich et al., 2006; Stitt et al., 2006) or leveraged sub-meter resolution or hyperspectral data (Lake et al., 2022; Parker Williams & Hunt, 2002), our study leveraged the deep temporal catalogue of (lower resolution) Landsat satellite imagery to detect leafy spurge over broad spatial scales with high detection accuracy (97%). Evaluating the model using ground-truth data from EDDMapS and iNaturalist indicated that our TempCNN model was effective across hundreds of thousands of square kilometers. Over the two decades of our study, models predicted a substantial increase in the area invaded (ca. 100,670 to 715,600 hectares).

These predictions are likely underestimates because we used a cautious threshold (p_leafy spurge_ = 0.5) that was close to the median probability calculated from our validation dataset (p_leafy spurge_ = 0.48). We chose that threshold to reduce false positives but acknowledge that this choice causes populations with a lower probability to be excluded from our calculations of invaded area (i.e. causes false negatives). However, we do not expect that leafy spurge classification errors affected the spatial and temporal trends predicted by our model. When we used a different probability threshold for leafy spurge (e.g. 0.38, which had 75% sensitivity for the validation dataset), spatial and temporal patterns of invasion were very similar to those reported for 0.5.

The general patterns that we observed also align with observations by researchers and land managers in this region and changes since previous estimates were reported (e.g. Skinner et al., 2002).

Although our TempCNN had high accuracy in detecting leafy spurge, the model still predicted false positives and false negatives as evidenced by moderate F2 and sensitivity scores, respectively. False positives may occur in areas that the model infers to be high-quality habitat (e.g. based on contextual information) but are not currently occupied or where the initial land classification map was incorrect (Stehman & Foody, 2019). False positives could also arise if our model confused leafy spurge with other plants that have a similar spectral profile. However, our model included both spectral and phenological information. Because our prior work has shown that phenology greatly improves specificity for leafy spurge (Lake et al., 2022), it is less likely that false positives are due to misidentification of plant species. Conversely, false negatives may occur because our model was less likely to predict small and low-density populations (Figure 2), likely due to decreased signal from fewer individuals. Future studies could reduce false negatives by incorporating higher spatial resolution imagery such that smaller patches of leafy spurge were more easily distinguishable from background vegetation. For example, one could combine Landsat and Sentinel-2 data products (i.e. Harmonized Landsat/Sentinel-2), which would increase the frequency of re-imaging (Pastick et al., 2020; Zhou et al., 2019). However, Sentinel-2 images are available only after 2015 and therefore were not feasible for estimating spatial and demographic trends in this study.

We found that leafy spurge was less apparent and challenging to detect during drought years, most likely because it was less likely to flower (Selleck et al., 1962). For example, in 2012-2014 when northwest Minnesota experienced severe drought, predicted leafy spurge area was reduced by 83% compared to the years immediately prior (2009-2011). After the drought ended (2015-2017), predicted leafy spurge area rebounded by 447% (Figure 3). While this biological response to drought affected detection, it also revealed that remote sensing could assess plant performance in relation to environmental variation. This finding also indicates that low detection probability during drought years may not necessarily equate to extirpation. Instead, multiple years of evidence are likely important for verifying eradication. Overall, time-series data help to buffer against making faulty inferences during unusual years, but environmental variation must still be accounted for given how strongly it can influence detection probability.

### Roadside association and management implications

Leafy spurge occurrences were more likely to be found near roadsides than random expectation for both remotely-sensed and community science datasets (Figure 5). The stronger roadside bias found in community science data is likely due to accessibility (Kadmon et al., 2004); this type of spatial bias can have important implications for the development of species distribution models because it influences the distribution of occurrence-environment relationships (Inman et al., 2021). The persistence of roadside bias in our remotely-sensed dataset indicates that roadsides are particularly suitable habitats, which has been observed for other invasive plants (Parendes & Jones, 2000; Tyser & Worley, 1992). In the context of management, our finding suggests that roads provide important pathways of invasion and that eradication along roads is critical for preventing future invasion.

### Remote sensing population dynamics

Understanding demographic variation across species’ ranges is important for determining the causes of range limits and factors modulating range shifts in response to climate change (Angert et al., 2011; Eckhart et al., 2011; Villellas et al., 2013). However, such datasets require enormous effort over long time periods and are not feasible for most systems. Our study suggests that remote sensing is a viable alternative in some systems for quantifying population dynamics and documenting range expansion/infilling. In this study, we used linear regression (e.g. slope coefficient) to characterize if populations were increasing, decreasing or lacked directional change. Positive slopes were concentrated in west-central grasslands and in the Twin Cities region; both regions also had the highest concentrations of recent occurrences in community science datasets (e.g. our 2021-2023 validation dataset). Whereas negative slopes were concentrated at the western edge of the range. In this area, biocontrol efforts have been common (Skinner et al., 2006). Thus, our inferences of population dynamics are consistent with independent observations of population growth or decline.

Our estimates of population change may have also been affected by factors unrelated to habitat suitability. Where leafy spurge has colonized new suitable habitat and occurs as small or low-density populations, we may have failed to detect occurrences in some years, especially during drought, and obtained erroneous demographic estimates. Although the variance in detection probability for small populations was relatively high, the median probability of detection was also fairly high, suggesting that many small populations were detected.

Agricultural conversion of grasslands to crops and invasive species management (e.g., herbicides, burning, and biocontrol applications) may have caused population declines unrelated to environmental conditions. While these influences undoubtedly occurred, they were infrequent and unlikely to affect the general temporal and spatial patterns that we report.

### Species distribution models affected by spatial bias

SDMs built using remotely-sensed data were less spatially biased than those built using community science datasets. Regardless of the model algorithm used, the remotely-sensed models had higher model discrimination (AUC) and were less overfit than the community science models. This regularization allowed for consistent prediction across geography, which was reflected in higher sensitivity in rural areas that were less well sampled in the community science datasets. For example, remotely-sensed SDMs predicted much higher habitat suitability in southeastern and western Minnesota compared to the community science models (Figures 7 & S10) . The community science models predicted highest suitability in primarily urban or disturbed (i.e. Interstate 35 in southern Minnesota) areas.

Incorporating demographic variation into species distribution models (SDMs) has been suggested to improve forecasts by providing more detailed information on where populations are growing, stable, or declining (Normand et al., 2014; Schurr et al., 2012). For the most part, slope-weighted models did not differ from traditional (or probability-weighted) models. One exception was that for Maxent models, slope-weighting improved discrimination (AUC) without causing overfitting. However, the projections of suitability did not differ substantially in terms of geographic extent. This result is not surprising given that the geography of occurrences is strongly correlated with the geography of positive slopes. In future studies with finer-grained time resolution, it may be possible to incorporate additional complexity (i.e. density-dependence) to refine inference of population dynamics. Such demographic insights may be particularly useful for improving species distribution models but may require caution as demography and occurrence probability may not be positively correlated (Thuiller et al., 2014).

### Conclusions

Our study expands remote sensing of an individual species to a large geographic scale and over a long time series despite modest resolution of imagery. Our approach also lays groundwork for future studies using higher-resolution and high-cadence satellite imagery (e.g., Sentinel, Planetscope), which are not currently available for long time series but will be in the future. We observed rapid population expansion (and some range expansion) that was concentrated in two disjunct regions near metropolitan areas that differ in climate. Remotely-sensed occurrences were less spatially biased than community science occurrences and produced more accurate species distribution models. However, the persistence of roadside bias in remotely sensed occurrences suggest that disturbance may be important to invasion and that roads may serve as invasion pathways. More comparisons of this sort are needed to evaluate the extent to which publicly-available occurrence datasets are spatially biased and the consequences for SDM predictions. Overall, examining the population dynamics of individual species over large geographic areas and long time series is feasible and should become more tractable as high-resolution imagery accumulates.

## ACKNOWLEDGEMENTS

We thank R. Venette for thoughtful discussions about analyses. We are grateful to the community scientists that collected occurrence data used in the study. All Landsat imagery was accessed through Google Earth Engine. The Minnesota Supercomputing Institute and Google Cloud provided computational and data storage resources.

## DATA ARCHIVING STATEMENT

Code, models, and sample data are available at: https://github.com/lake-thomas/leafy-spurge-demography. Raster predictions are archived and available through DOI: XXX (UMN DRUM – will be uploaded at publication acceptance)

## AUTHOR CONTRIBUTIONS

TAL, RBR, and DM conceived of and designed the research study. TAL collected data for and developed remote sensing models. RBR collected data for and developed species distribution models. TAL and RBR performed the analyses. TAL, RBR, and DM contributed substantially to the drafting, editing, and final version of the paper.

