## Supplementary Materials for "Two decades of satellite images reveal the spatial and temporal dynamics of leafy spurge invasion and improve species distribution models"

[1]Co-first authors

[6]https://orcid.org/0000-0002-8836-5164

[7]https://orcid.org/0000-0001-7160-9110

[8]https://orcid.org/0000-0002-6202-9912

**Table of Contents:**

Supplementary Methods
SM 1.1 Landsat satellite imagery
SM 1.2 Ground truth maps
SM 1.3 TempCNN training data
SM 1.4 Temporal convolutional neural network (TempCNN) architecture
SM 1.5. TempCNN model training
SM 1.6 TempCNN model evaluation
SM 1.7 TempCNN model predictions
SM 1.8 Assessing spatial bias near roadsides

Supplementary Figures
Figure S1. Training history of the temporal convolutional neural network.
Figure S2. Occurrence points and polygon records of leafy spurge in Minnesota.
Figure S3. Thresholding predictions for leafy spurge occurrences.
Figure S4. Bioclimatic variables used for species distribution models.
Figure S5. Cross-validation spatial blocks for species distribution models.
Figure S6. Remote sensing predictions of leafy spurge at less than 5% probability used to generate absences for BRT models.
Figure S7. Independent records of leafy spurge from 2021-2023 partitioned by urban and non-urban regions.
Figure S8. Normalized confusion matrix displaying classification results of the temporal convolutional neural network for 15 land cover classes and leafy spurge.
Figure S9. Histograms of predicted leafy spurge and land cover class probabilities (2018-2020) sampled against the 2021 NLCD dataset.
Figure S10. Boosted regression tree (BRT) model projections of climate suitability.

Supplementary Tables
Table S1. Community science occurrence data of leafy spurge from 2000 to 2023.
Table S2. Number of training, validation, and testing records of Landsat satellite pixels for training the temporal convolutional neural network.
Table S3. Temporal convolutional neural network model performance and evaluation metrics. Model performance for leafy spurge and 15 additional land cover classes are based on the number of true positives (TP), false positives (FP), true negatives (TN) and false negatives (FN) in the testing dataset.
Table S4. Area of leafy spurge invasion in square kilometers predicted from the TempCNN model between 2000 and 2020.
Table S5. Tests for differences in the extent of leafy spurge invasion across different geographic regions and mean Palmer Drought Severity Index (PDSI) values.

Supplementary References

### Supplementary Methods

*SM 1.1 Landsat satellite imagery*

We obtained Landsat 5 TM, 7 ETM+, and 8 OLI surface reflectance products (Collection 2 Tier 1) from 2000 to 2020 via the Google Earth Engine platform. We began by removing pixels with cloud, cloud shadow, and radiometric saturation using quality assessment bitmasks ('QA_Pixel' and 'QA_Radsat'). After cleaning the data, we created image mosaics for two-month periods by calculating the median pixel value for each spectral band. This process yielded a Landsat time-series dataset composed of seasonal image mosaics from 2000 to 2020, one from each of the following two-month periods: March-April, May-June, and July-August.

For instance, if six Landsat images captured between March and April in one year have pixel values for the 'blue' band as (0.51, 0.55, 0.62, NA, 0.62, 0.73), the NA values indicate data removed during quality assessment. We then calculated the median value for each spectral band and pixel to construct the seasonal mosaic using the 'ee.ImageCollection.median' function in Google Earth Engine. We repeated this process to create mosaics for March-April, May-June, and July-August across all available images for each year from 2000 to 2020.

*SM 1.2 Ground truth maps*

We used the National Land Cover Database (NLCD) to associate Landsat pixels with land cover classes for classification. The NLCD provides a 30-meter resolution classification of 20 land cover classes across the conterminous US for nine years (2001, 2004, 2006, 2008, 2011, 2013, 2016, 2019, and 2021). We used pixel-based reclassification to include leafy spurge occurrence points (N = 3,345) into the existing NLCD maps based on reported survey dates (Table S1). For example, leafy spurge occurrences reported in 2018, 2019, and 2020 were added to the nearest NLCD map (2019) by reclassifying existing land cover types.

The 15 NLCD land cover classes present in Minnesota include: Open Water, Developed Open Space, Developed Low Intensity, Developed Medium Intensity, Developed High Intensity, Barren Land, Deciduous Forest, Evergreen Forest, Mixed Forest, Shrub/Scrub, Grasslands/Herbaceous, Pasture/Hay, Cultivated Crops, Woody Wetlands, and Emergent Herbaceous Wetlands.

*SM 1.3 TempCNN training data*

Neural networks benefit from large, representative datasets to learn to classify input data to corresponding labels. We randomly sampled one hundred thousand pixels across Minnesota. At each sampled location, we extracted Landsat spectral data—spanning the periods of March/April, May/June, and July/August—across three consecutive years, along with the corresponding latitude, longitude, and land cover class for an NLCD year (Figure 1). For example, in the 2019 NLCD year, we obtained Landsat data from 2018, 2019, and 2020, along with the latitude, longitude, and land cover class from the NLCD map for one hundred thousand samples. We repeated this process for seven NLCD maps yielding approximately seven hundred thousand samples (Tables S1 & S2).

*SM 1.4 Temporal convolutional neural network (TempCNN) architecture*

We developed a supervised deep learning model (TempCNN) to classify leafy spurge and 15 additional land cover classes. The TempCNN architecture has shown to produce higher classification accuracy compared to shallower machine learning algorithms (e.g., Random Forests) for satellite image time series classification because it can directly learn from the temporal dimension of the data (Allred et al. 2021, Brock and Abdallah, 2022). The neural network contains a series of convolutional layers (1D convolutions), dropout layers, and fully connected layers.

In deep learning, convolutions are mathematical operations that are typically applied to images. Familiar examples of convolutions that are applied to images are "blur" or "sharpening" filters. Here, we use 1D convolutions to transform our inputs (a vector of spectral data from nine seasonal mosaics in our Landsat time-series) (i.e., March/April, May/June, and July/August for the three consecutive years) to learn the association between spectral and phenological signals and land cover classes. Specifically, a 1D convolution is a small filter that looks at a subset of the data and performs a mathematical transformation on the grouping of data to output new synthetic higher order information about the data group. The size of the convolution filter (i.e., the kernel size) can also vary to allow the model to learn from different sets of spectral data and wider phenological windows. The convolution then steps (or, slides) over the data to perform the same operation on the next group. The deep learning algorithm uses training to learn parameter values for the transformations of the convolutions that lead to the most accurate classification.

Deep learning models are subject to overfitting because of the large number of parameters. We used dropout layers to minimize the problem of overfitting. These layers randomly set a selection of parameters in the model to zero as a form of regularization to prevent the model from memorizing the training data. By randomly deactivating certain parameters during the training process, dropout encourages the network to learn in a more robust and generalizable manner.

Fully connected layers connect all parameters from previous layers to allow for learning between layers. Additionally, we included a concatenate layer to combine spectral and phenological features with location data (i.e., latitude, longitude) to provide spatial context during training. Overall, the model outputs a vector of probabilities for each land class for each pixel (range: 0-1). Finally, to generate pixel classifications to test model accuracy, we used an Argmax layer that selected the land cover class with the highest probability as the land class assignment. Additional details on the model architecture are available through a GitHub repository: https://github.com/lake-thomas/leafy-spurge-demography/

*SM 1.5. TempCNN model training*

We randomly divided the seven hundred thousand samples into an 80/10/10 ratio of training, validation, and testing data (Table S2). The validation data is used to evaluate and fine-tune the model's parameters during training. The testing data is withheld from model training and used as an independent set to evaluate performance.

Training the deep learning model comprises of two main steps - the forward pass and the backward pass.

Initially, the model's parameters (i.e., weights and biases) were randomly selected from a uniform distribution. During the forward pass, samples of the input data (in this case, 8,192 samples of spectral data from our Landsat time-series) are fed through the model. As the data moves through each layer of the network, various computations are performed through a series of convolutions and dropout layers using the model's current parameters. The goal is to produce an output that is as close as possible to the actual target output (the land cover class). The discrepancy between the predicted output and the actual land cover class is calculated as the training loss using a loss function. We used the categorical cross-entropy loss to measure the difference between predicted labels and true labels for each class, as is common in multi-class classification problems.

Once the loss is computed from the training data, the optimizer determines how to adjust the model's parameters to improve its performance. This is done during the backwards pass. During this stage, the model calculates gradients, which are essentially directions and magnitudes of change needed for each parameter to minimize the loss. The Adam optimizer (Kingma & Ba, 2014), a variant of stochastic gradient descent, uses these gradients to update the model's parameters. Adam is popular because it incorporates adaptive learning rates and momentum, making the optimization process more efficient by speeding up convergence and stabilizing updates.

By repeating the forward and backward passes over multiple epochs—complete passes over the training dataset—the model progressively learns the intricate patterns in the data, allowing it to make accurate predictions when presented with new, unseen testing data. Throughout the process, the validation dataset helps in tuning model parameters to prevent overfitting. After each epoch, the loss is also computed against the entire validation dataset (i.e., the validation loss). Validation loss is crucial because it helps assess how well the model generalizes to new, unseen data. Ideally, the validation loss should also decrease as training progresses. However, if it starts to increase significantly while the training loss continues to decrease, this indicates that the model might be overfitting to the training data. As such, we trained the TempCNN for 100 epochs and monitored both the training and validation loss to prevent overfitting (Figure S1).

Due to the uneven number of samples per land cover class, we computed a class weight vector as the inverse frequency of class counts in the training dataset. Class weights were applied to the loss function during the training process to increase the importance of errors made on rare classes relative to more common classes (Cabezas et al., 2020) (Table S2).

We set the dropout rate at 10% (Brock & Abdallah, 2022). For convolutional layers, we used a kernel size of 3 and increased the number of kernels from 64 to 128 after the first layer. We used dilated convolutions with dilation rates of 1, 2, and 4 for each successive layer. Dilated convolutions have modified kernels with spaces (or, gaps) between each parameter. These gaps effectively increase the size of the kernel without adding additional parameters to the model. The dilated convolutions also increase the model's receptive field to capture both annual and inter-annual phenological signals (Pelletier et al., 2019).

We trained the TempCNN using an adaptive learning rate for the Adam optimizer. Specifically, we used learning rate warmup and cosine decay. Initially set at 1e-7, the learning rate linearly increased to 1e-2 over the first 10 epochs, after which it gradually decayed in a cosine pattern for the remaining epochs. This strategy promotes stable initial training with a slow learning rate and efficient convergence to improve performance and generalization. We trained the TempCNN using Python v.3.7.1 with Tensorflow v.2.0.0 (Abadi et al., 2016) and Keras v.2.10.0 (Chollet, 2017).

*SM 1.6 TempCNN model evaluation*

We assessed model performance for each class for several evaluation metrics that are based on a class-wise confusion matrix from the Argmax output. The confusion matrix contains the number of correctly predicted true positives (TP), incorrectly predicted false positives (FP), correctly predicted true negatives (TN), and incorrectly predicted false negatives (FN) for all records in the withheld testing dataset.

The evaluation metrics we calculated were: Accuracy, Specificity, Sensitivity, Precision, and F2.

We calculated overall accuracy as the proportion of correctly identified pixels:

Accuracy = (TP + TN) / (TP + TN + FP + FN).

We calculated class-wise specificity, or the ability to discriminate true negatives (i.e. True Negative Rate (TNR)).

TNR = TN / (TN + FP).

We calculated class-wise sensitivity (also termed recall), or the ability to predict all true positives (i.e. True Positive Rate (TPR)).

TPR = TP / (TP + FN).

We calculated the F2-score per class. F2 is derived from precision (i.e. the proportion of all identified positives that are correct) and sensitivity (i.e. TPR, see above). F-scores are employed when both precision and sensitivity are important factors for model evaluation. F2 places more importance on sensitivity relative to precision. This is preferred as a performance metric for invasive species because the metric weighs false negatives greater than false positives due to the fact that the potential economic cost of falsely declaring an invasive species absent is high.

Precision = TP / (TP + FP).

F2-score = [(5 * precision * sensitivity) / (4* (precision + sensitivity)].

*SM 1.7 TempCNN model predictions*

We predicted the probability of leafy spurge and 15 additional land cover classes at a 30-meter resolution across Minnesota, totaling approximately 250 million pixels. The probability of each land cover class can be interpreted as the likelihood that the class is present in each pixel. As the TempCNN required the same data structure for prediction as used for training (i.e., three consecutive years of Landsat data), we predicted each land cover class across seven time-steps: 2000-2002, 2003-2005, 2006-2008, 2009-2011, 2012-2014, 2015-2017, and 2018-2020.

To gauge the predictive capabilities of our TempCNN across land cover classes, we evaluated the model against the NLCD's 2021 dataset. For each land cover class, we randomly sampled 10,000 points from the NLCD 2021 layer, extracted the predicted probability values at these locations from our most recent TempCNN model predictions (2018-2020), and plotted histograms to visualize agreement between our predictions and the NLCD dataset (Figure S9).

To further gauge the predictive capabilities of our TempCNN for leafy spurge, we evaluated performance on an independent set of occurrence points (N = 1,018) and polygons (N = 940) with reported population area estimates ('Infested Area, $m^2$') collected from iNaturalist and EddMaps between the years 2021-2023 (Figure S2).

We also examined how well the latest leafy spurge prediction (2018-2020) captures newly reported occurrences in 2021-2023. For each occurrence in 2021-2023, we extracted the predicted probability from the 2018-2020 leafy spurge model prediction and plotted these probabilities against the number of occurrence records (Figure S9).

To assess the models' ability to detect populations of different sizes, we used polygons with reported population area estimates (measured as 'Infested Area, $m^2$'). Because the polygons varied in size, we averaged predicted probabilities across all pixels intersecting each polygon. This approach ensured that we captured the variation in density and spatial heterogeneity within populations. We then fit quantile regressions between the mean predicted probability of leafy spurge and population area for a range quantiles to determine how population size influences the likelihood of leafy spurge occurrence using the 'rg' function from the 'quantreg' package (Koenker et al., 2018). To explore how the intercepts and slopes changed across all quantiles, we first fit the model with tau = -1, which explores quantiles (0,1). We then fit a representative sample to report with tau = c(0.05, 0.25, 0.5, 0.75) (tau = 0.5 represents regression on the median). This allowed us to examine whether the 2018-2020 model was more or less sensitive to detecting smaller infestations relative to larger ones (Figure 2).

*SM 1.8 Assessing spatial bias near roadsides*

We measured the distance to primary and secondary roads for both community science and remotely-sensed occurrence points using the U.S. Census Bureau's MAF/TIGER Database shapefiles for roads in Minnesota. We used the same occurrence points from the SDM analyses (N = 1,771 occurrences per dataset). We then used a k-d tree nearest neighbor search to calculate the shortest distance between occurrence points and road segments. For comparison, 1,771 random points were sampled from within the Minnesota state boundary and distances to roads were similarly calculated. We repeated the random point sampling process for 999 permutations and calculated the mean distance to roads. We evaluated spatial bias by comparing the mean distance of community science, remote sensing, and random observations to roadsides (Figure 4).

**Supplementary Figures**

Figure S1. Training history of the temporal convolutional neural network (TempCNN). The left panel shows the training (blue) and validation (orange) loss across 100 epochs, while the right panel displays training and validation accuracy over the same period. The training loss and accuracy steadily improve and converge.

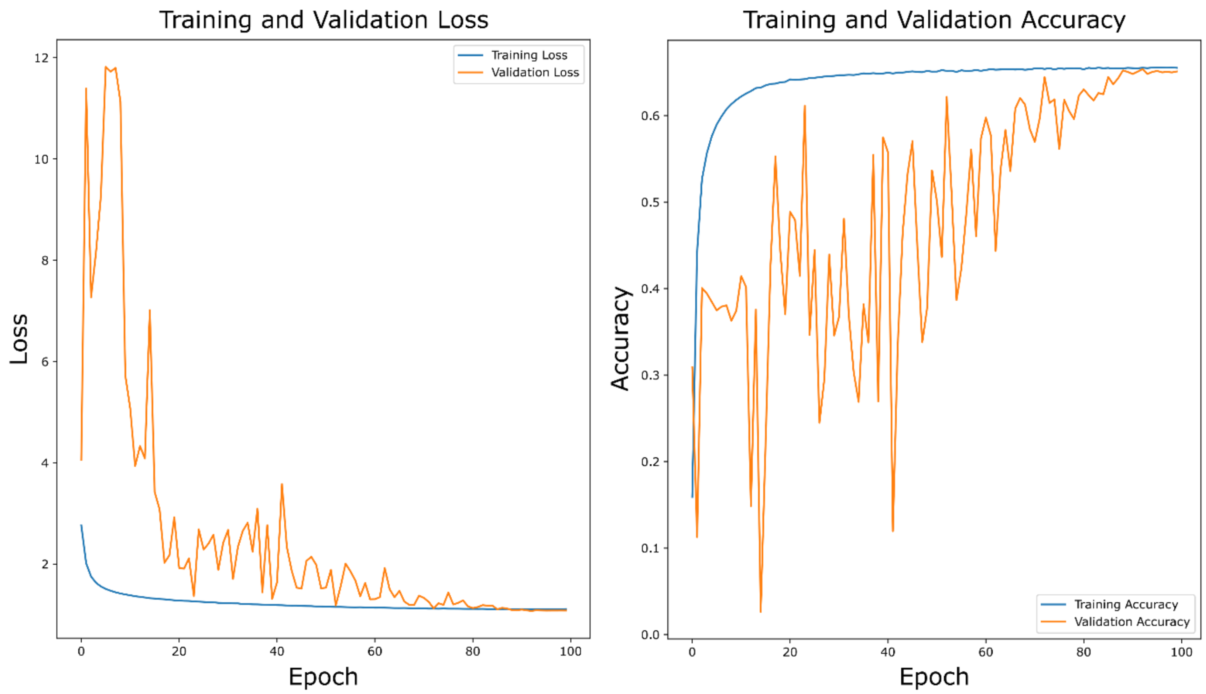

Figure S2. Occurrence points and polygon records of leafy spurge in Minnesota. The left panel shows occurrence points (blue) from 2000-2020 (N=3,345). The middle panel shows occurrence points (red) from 2021-2023 used for TempCNN model evaluation (N=1,018). The right panel shows polygon centroids (green) with reported population size estimates from 2021-2023 used to evaluate the TempCNN model (N=940).

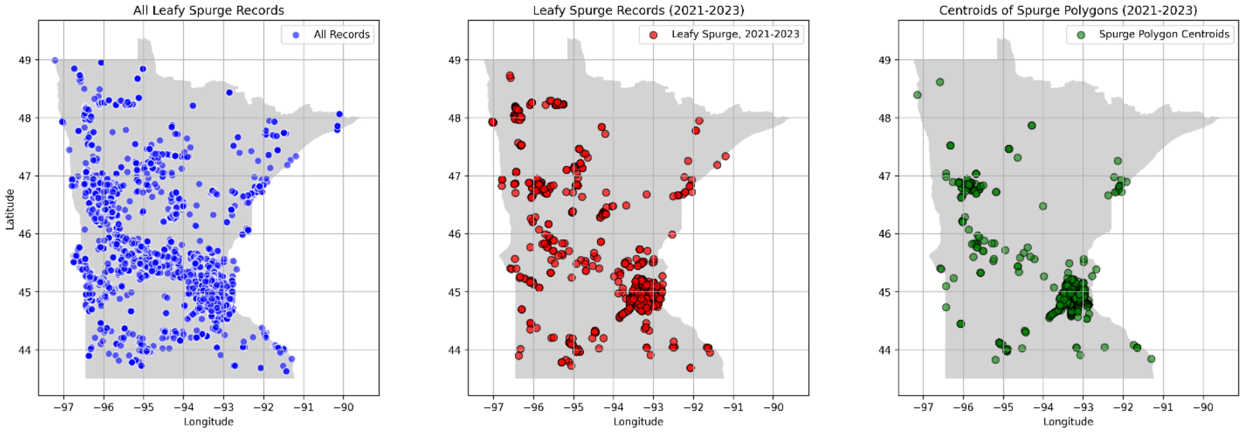

Figure S3. We transformed TempCNN predictions of leafy spurge into discrete occurrence points. We chose a probability threshold for a positive leafy spurge presence of 0.5. We chose this threshold because it indicated when the probability of leafy spurge was greater than the sum of the probability of all other classes. It may be taken as a conservative threshold estimate because it was greater than the that corresponded to threshold required to obtain 75% sensitivity of occurrences reported by community scientists using the 2009-2011 prediction period as a midpoint (threshold = 0.38 estimated at the 2009-2011 midpoint prediction period).

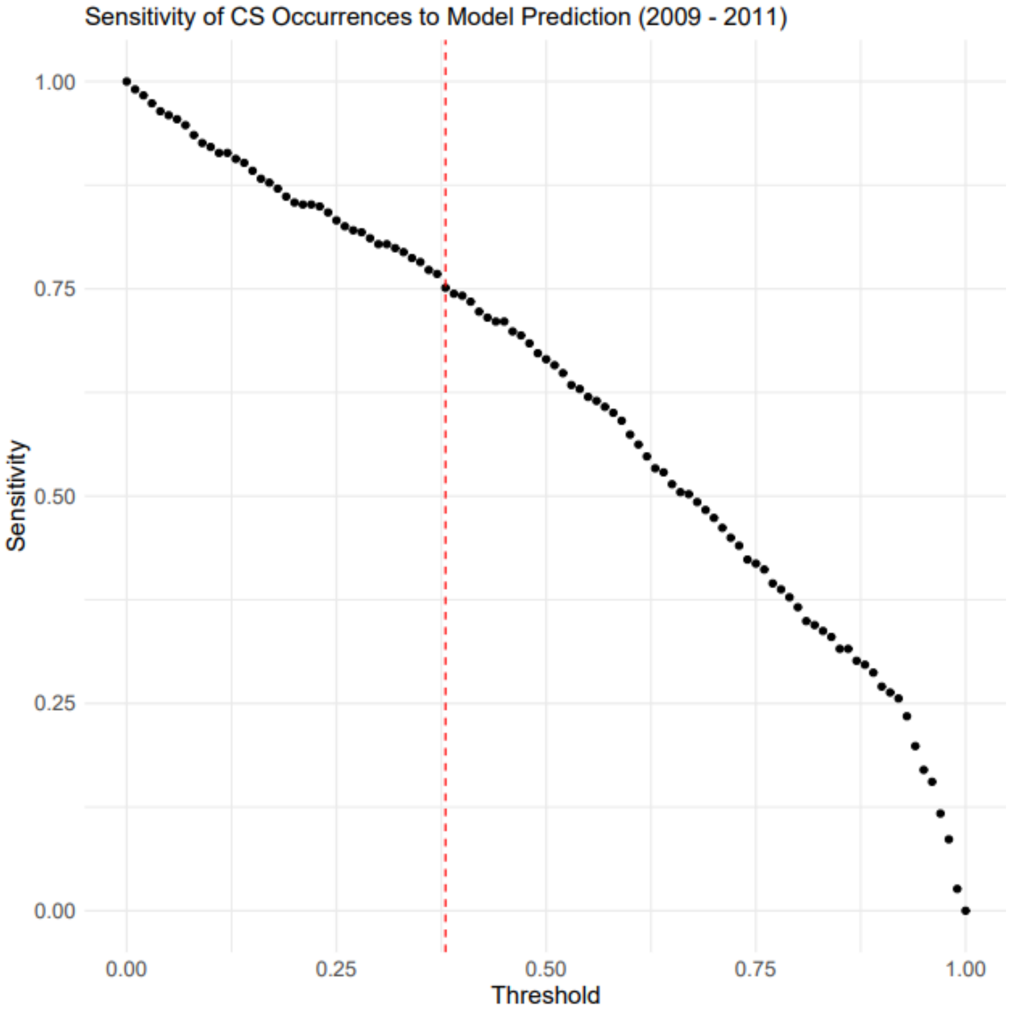

Figure S4. Bioclimatic variables used for species distribution modelling, including mean temperature of the warmest quarter (Bio 10), mean temperature of the coldest quarter (Bio 11), and annual precipitation (Bio 12).

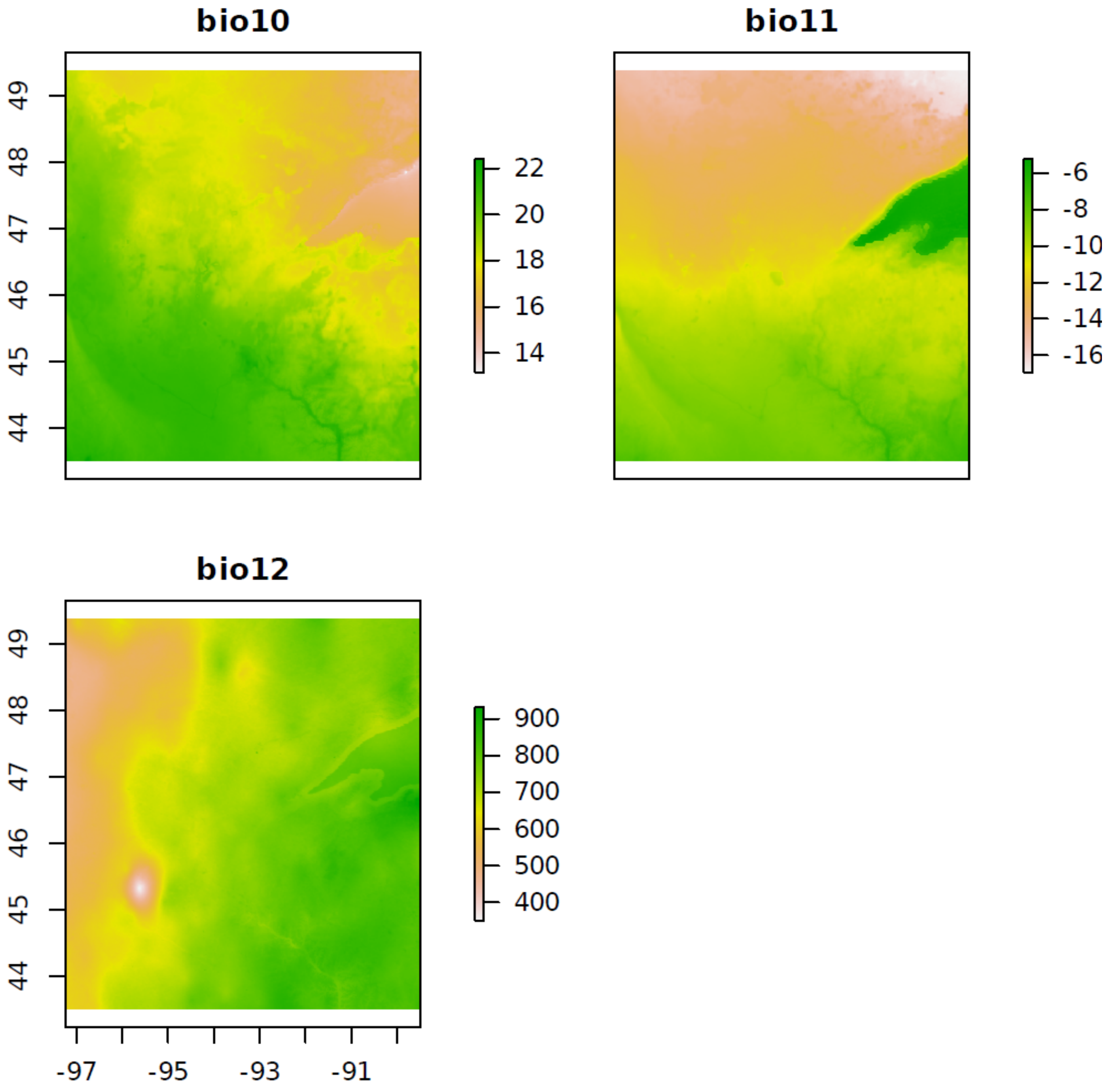

Figure S5. Cross-validation spatial blocks for species distribution models using community science and remote sensing occurrences datasets.

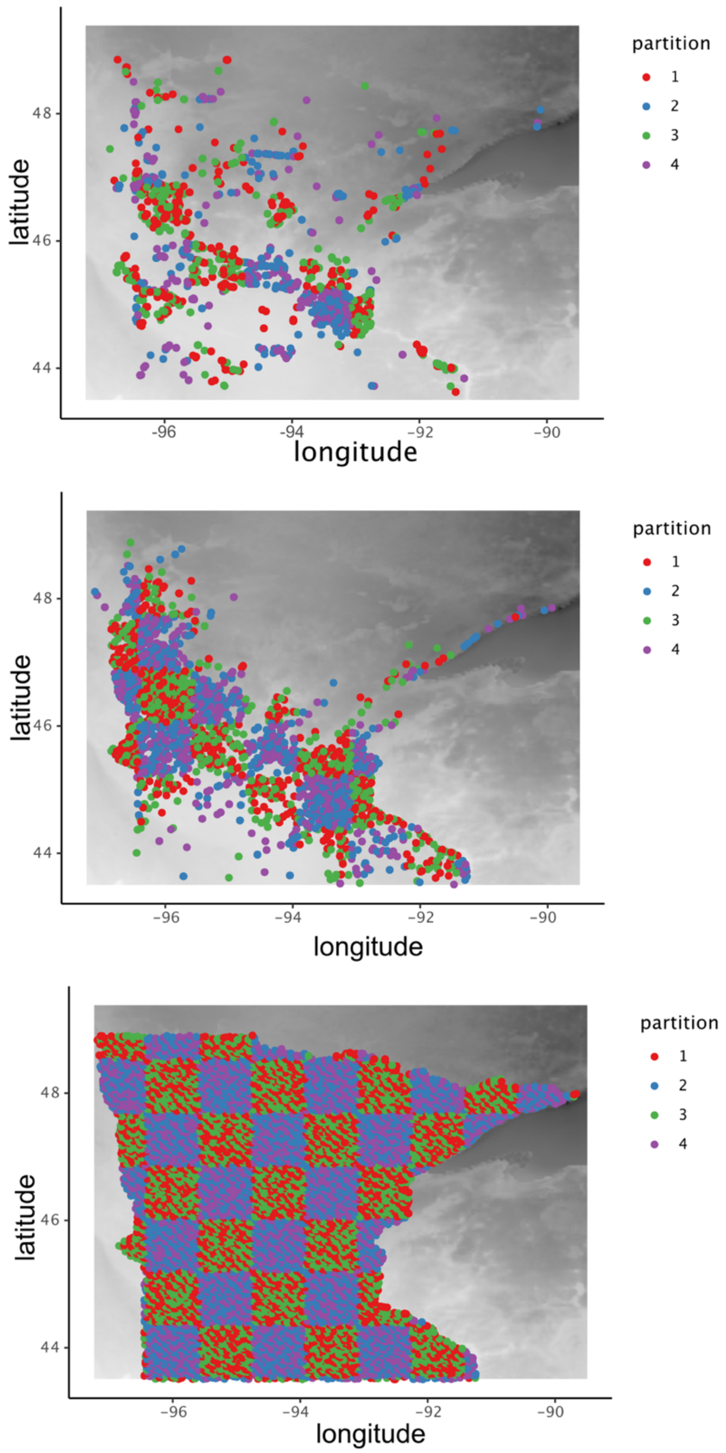

Figure S6. Binary map of the predicted probability of leafy spurge with areas < 0.05 probability shown in green, and areas > 0.05 probability shown in white. Absence points were sampled in areas with a < 0.05 probability for the Boosted Regression Tree (BRT) models.

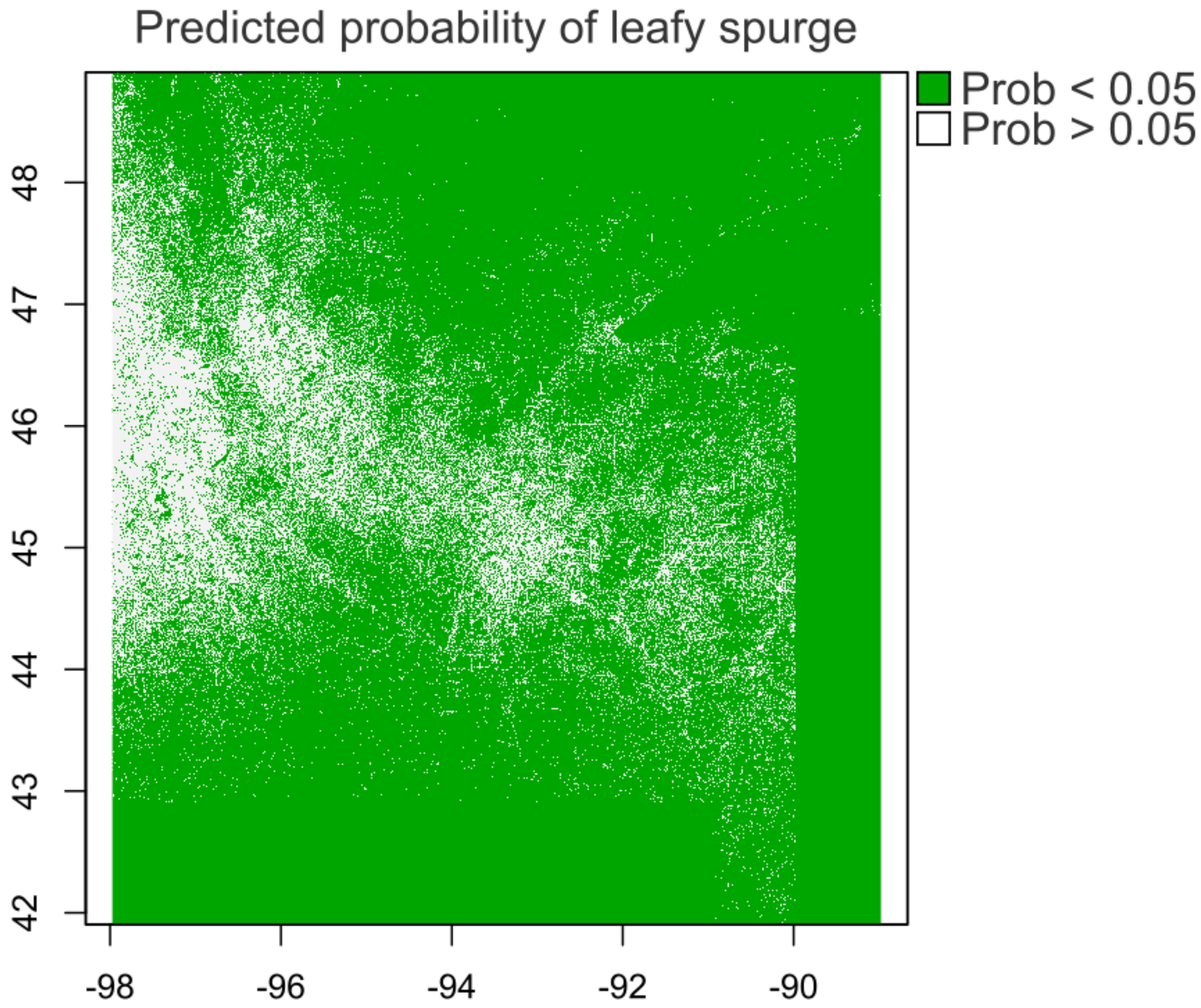

Figure S7. Independent records of leafy spurge from 2021-2023 partitioned by geographic region. Red squares indicate records sampled from the surrounding metropolitan region of Minnesota (including the cities of Minneapolis and St. Paul, MN). Blue circles indicate records sampled outside of the metropolitan center. When evaluating SDMs, we calculated sensitivity separately for all points and non-metropolitan points (blue) to assess model discrimination in different geographic contexts.

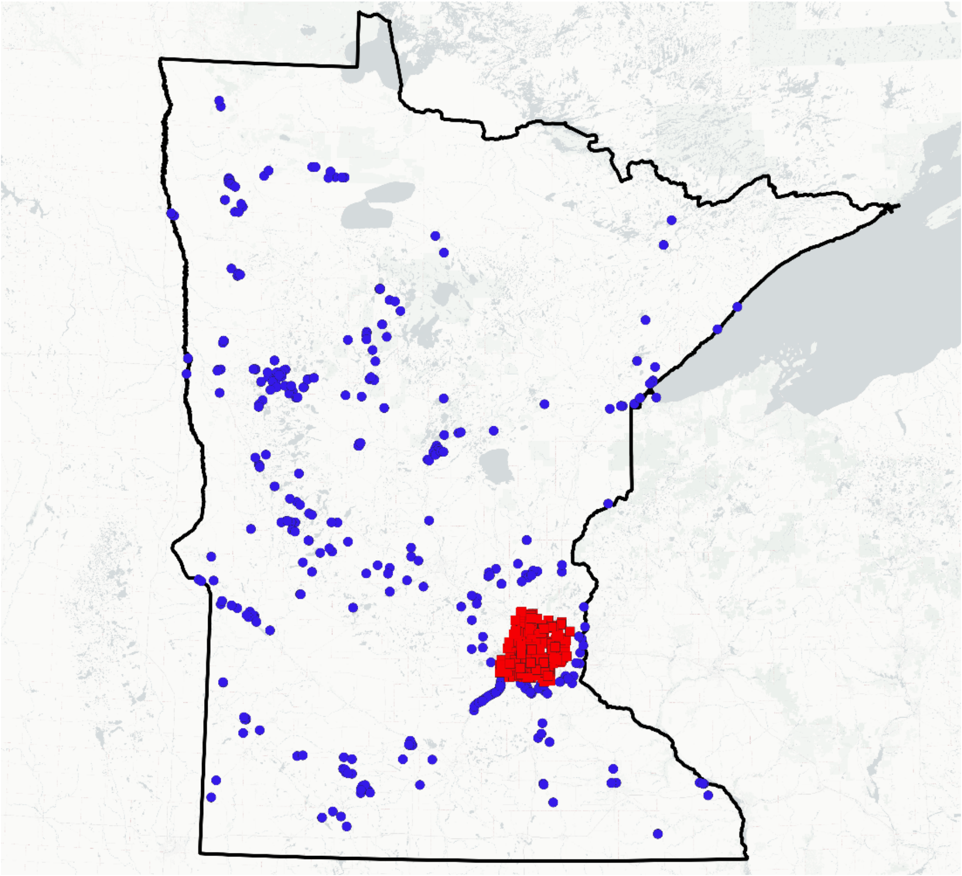

Figure S8. Confusion matrix displaying classification results of the temporal convolutional neural network based on the 10% withheld testing dataset. Classification results were calculated from the testing dataset for 15 land cover classes and leafy spurge. Values in the confusion matrix are normalized along rows, where the diagonal represents the proportion of records for which the predicted class is equal to the true class (i.e., true positive rate). Off-diagonal elements are misclassifications between the true class and the predicted class.

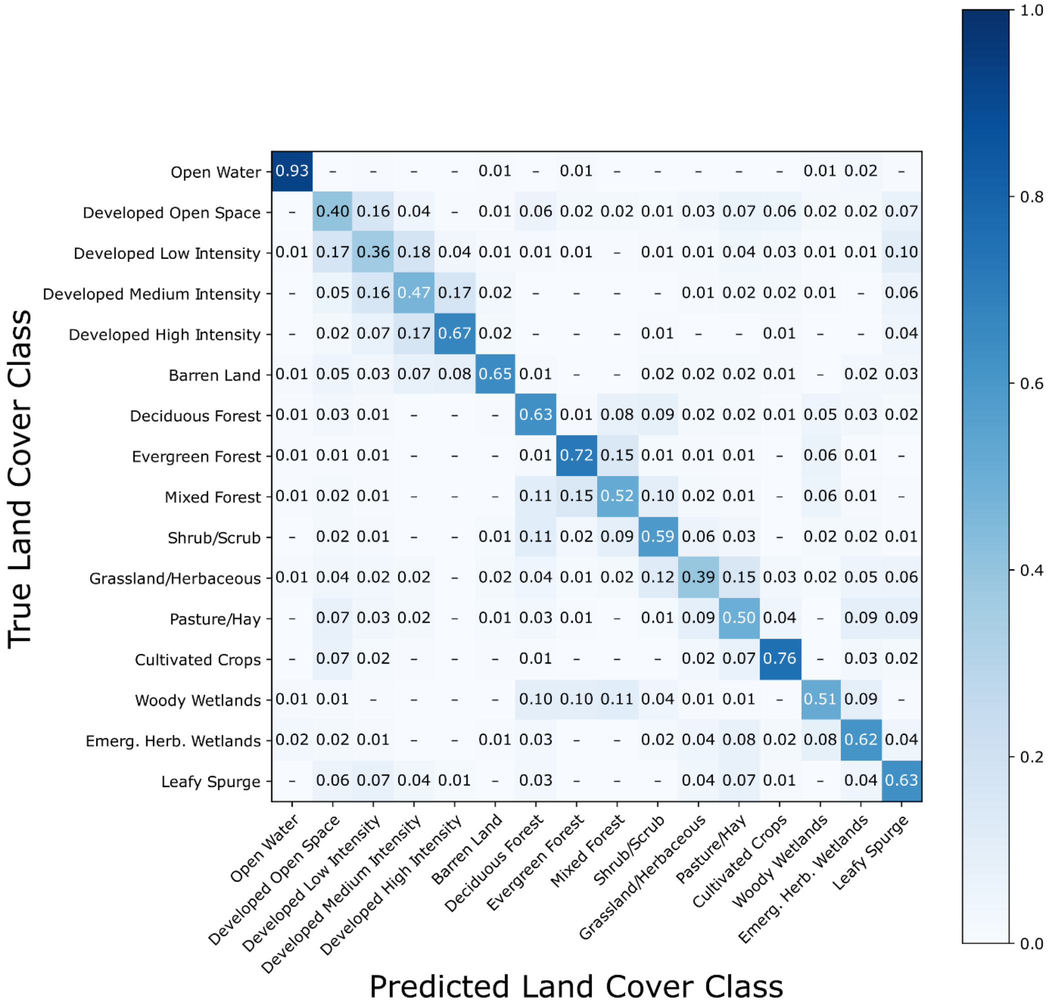

Figure S9. Comparison of predicted land cover class probabilities from our latest prediction (2018-2020) with the National Land Cover Database's 2021 dataset. Histograms of predicted probabilities for 15 predicted land cover classes (purple histograms) were created by randomly sampling 10,000 points per land cover class from the NLCD 2021 dataset and extracting the probability values for these points from our latest predictions. In addition, we generated a leafy spurge prediction across Minnesota using Landsat imagery from 2018-2020 and determined if recently reported occurrences (from 2021-2023, N = 1,018) were identified with high probability by plotting the histogram of probabilities (red histogram). For each plot, the Y-axis indicates the number of random land cover class samples. The X-axis indicates the probability of each land cover class predicted in 2018-2020. The dashed vertical line indicates the median probability for each histogram.

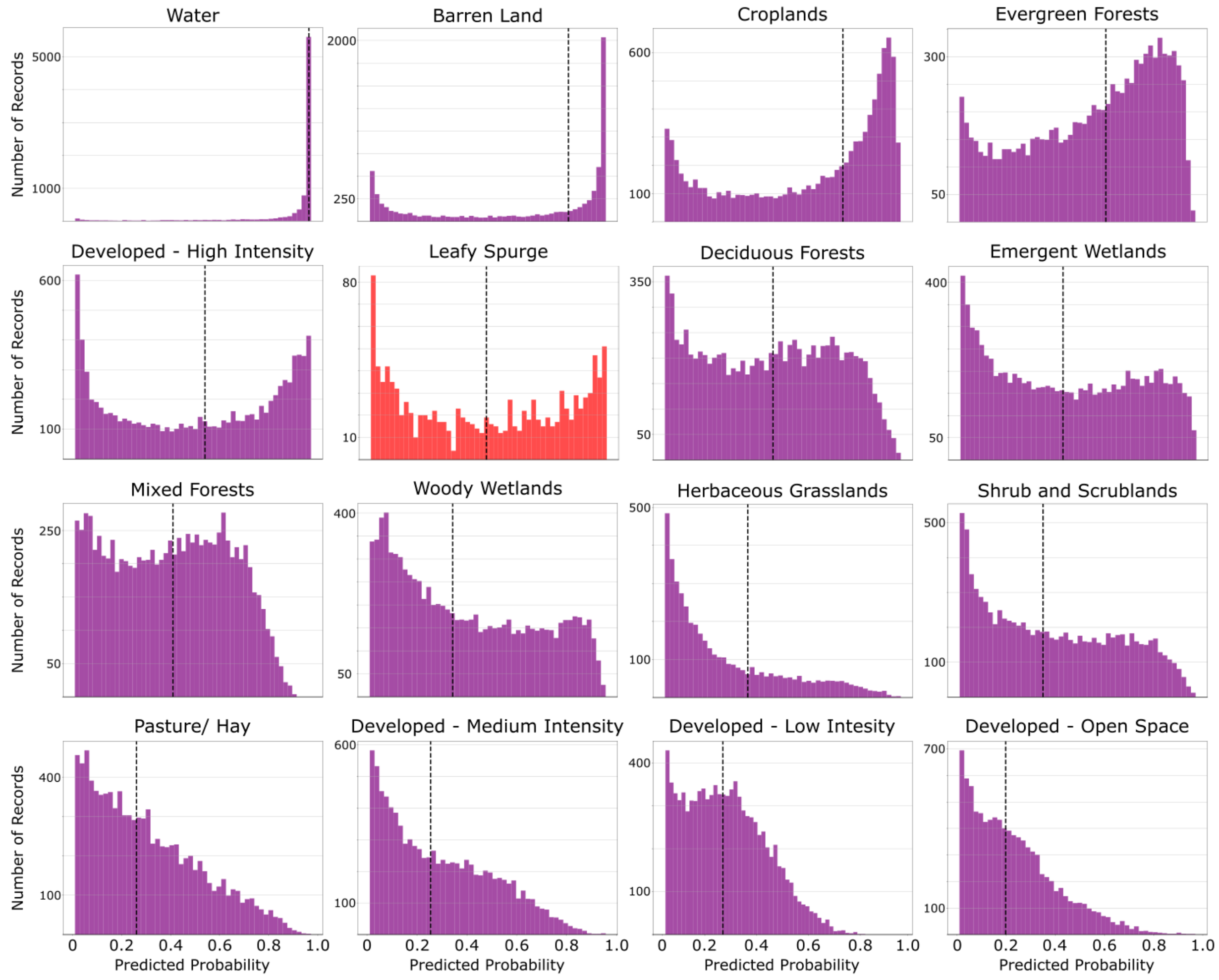

Figure S10. Boosted regression tree (BRT) model projections of climate suitability from models built using either community science or remotely-sensed occurrence datasets and weighting systems (traditional, probability, and slope). Each model projection is the average of nine models built with different combinations of hyperparameters (see text for more details).

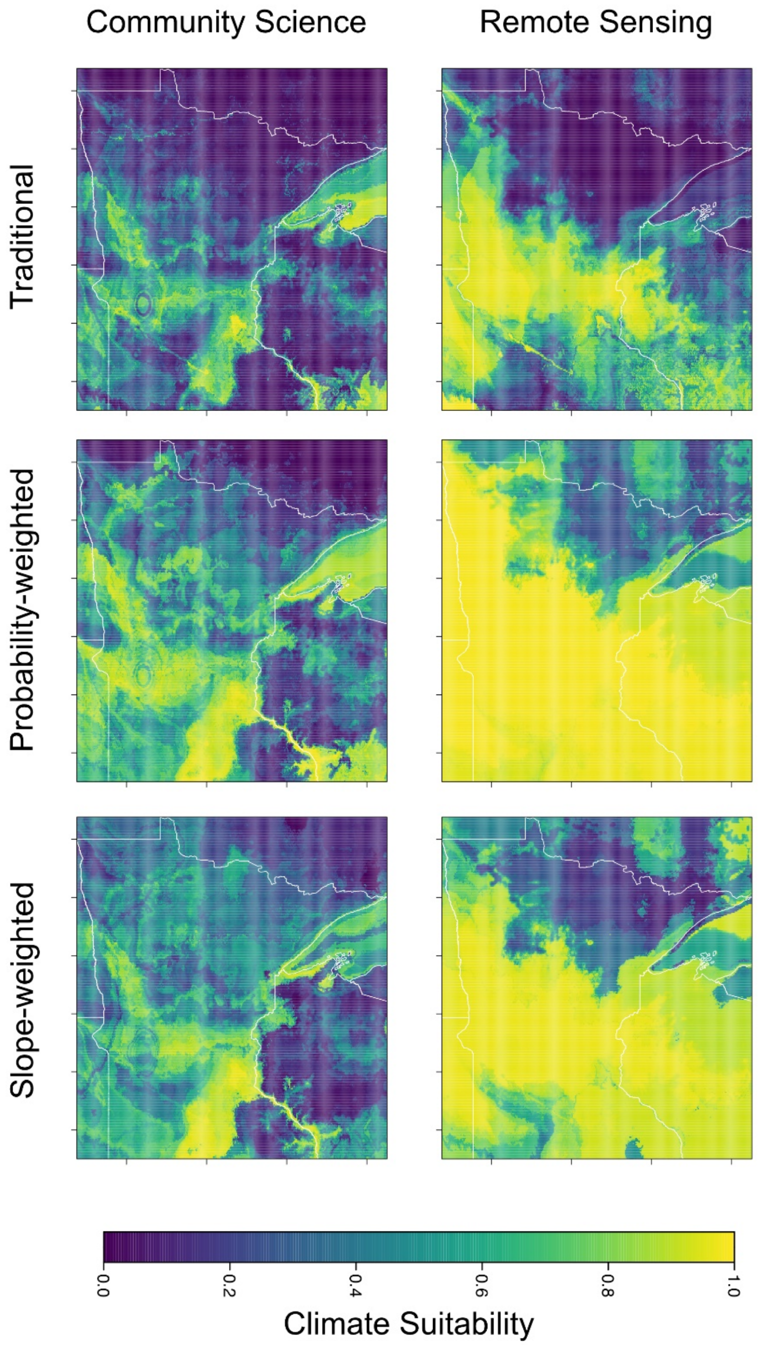

### Supplementary Tables

Table S1. Community science occurrence data of leafy spurge from 2000 to 2023.

| Year | EddMaps | iNaturalist | Total | NLCD Year |
|---|---|---|---|---|
| 2000 | 31 | 0 | 31 | 2001 |
| 2001 | 26 | 0 | 26 | |
| 2002 | 6 | 0 | 6 | |
| 2003 | 4 | 0 | 4 | 2004 |
| 2004 | 96 | 0 | 96 | |
| 2005 | 35 | 0 | 35 | |
| 2006 | 387 | 0 | 387 | 2008 |
| 2007 | 49 | 0 | 49 | |
| 2008 | 183 | 1 | 184 | |
| 2009 | 140 | 0 | 140 | 2011 |
| 2010 | 175 | 0 | 175 | |
| 2011 | 80 | 1 | 81 | |
| 2012 | 122 | 0 | 122 | 2013 |
| 2013 | 35 | 3 | 38 | |
| 2014 | 30 | 1 | 31 | |
| 2015 | 44 | 2 | 46 | 2016 |
| 2016 | 116 | 1 | 117 | |
| 2017 | 68 | 9 | 77 | |
| 2018 | 100 | 20 | 120 | 2019 |
| 2019 | 568 | 73 | 641 | |
| 2020 | 862 | 77 | 939 | |
| 2021 | 383 | 89 | 472 | N/A |
| 2022 | 315 | 98 | 413 | |
| 2023 | 73 | 60 | 133 | |
| Total Leafy Spurge Occurrences (2000-2020) | | | 3345 | |

Table S2. Number of training, validation, and testing records of Landsat satellite pixels for each land cover class and leafy spurge occurrences. Corresponding class weights calculated as the inverse proportion of the number of records in the training data for training the temporal convolutional neural network.

| Class | Training Samples | Testing Samples | Validation Samples | Class weight |
|---|---|---|---|---|
| Cultivated Crops | 214134 | 26734 | 26466 | 2.6 |
| Woody Wetlands | 86666 | 10820 | 10712 | 6.4 |
| Deciduous Forest | 62000 | 7741 | 7663 | 9.0 |
| Emergent Herbaceous Wetlands | 46052 | 5749 | 5691 | 12.1 |
| Pasture/ Hay | 32310 | 4033 | 3993 | 17.2 |
| Open Water | 30184 | 3768 | 3731 | 18.4 |
| Mixed Forest | 27185 | 3394 | 3360 | 20.4 |
| Developed Open Space | 14700 | 1835 | 1817 | 37.8 |
| Evergreen Forest | 12626 | 1576 | 1560 | 44.0 |
| Developed Low Intensity | 9142 | 1141 | 1130 | 60.8 |
| Grassland/ Herbaceous | 6794 | 848 | 840 | 81.8 |
| Shrub/ Scrub | 4583 | 572 | 566 | 121.3 |
| Developed Medium Intensity | 4422 | 552 | 547 | 125.7 |
| Leafy Spurge | 2680 | 335 | 330 | 207.4 |
| Developed High Intensity | 1403 | 175 | 173 | 396.2 |
| Barren Land | 1012 | 127 | 125 | 549.3 |

Table S3. Temporal convolutional neural network model performance and evaluation metrics. Model performance for leafy spurge and 15 additional land cover classes are based on the number of true positives (TP), false positives (FP), true negatives (TN) and false negatives (FN) in the testing dataset.

| Class Name | TP | TN | FP | FN | Accuracy | Sensitivity | Specificity | Precision | F2 |
|---|---|---|---|---|---|---|---|---|---|
| Leafy Spurge | 217 | 67223 | 1842 | 118 | 0.97 | 0.65 | 0.97 | 0.11 | 0.32 |
| Open Water | 3506 | 65256 | 376 | 262 | 0.99 | 0.93 | 0.99 | 0.9 | 0.92 |
| Developed Open Space | 714 | 64477 | 3088 | 1121 | 0.94 | 0.39 | 0.95 | 0.19 | 0.32 |
| Developed Low Intensity | 414 | 67086 | 1173 | 727 | 0.97 | 0.36 | 0.98 | 0.26 | 0.34 |
| Developed Medium Intensity | 260 | 68282 | 566 | 292 | 0.99 | 0.47 | 0.99 | 0.31 | 0.43 |
| Developed High Intensity | 119 | 69010 | 215 | 56 | 0.99 | 0.68 | 0.99 | 0.36 | 0.58 |
| Barren Land | 89 | 68848 | 425 | 38 | 0.99 | 0.7 | 0.99 | 0.17 | 0.44 |
| Deciduous Forest | 4927 | 59499 | 2160 | 2814 | 0.93 | 0.64 | 0.96 | 0.7 | 0.65 |
| Evergreen Forest | 1136 | 66094 | 1730 | 440 | 0.97 | 0.72 | 0.97 | 0.4 | 0.62 |
| Mixed Forest | 1786 | 63872 | 2134 | 1608 | 0.95 | 0.53 | 0.97 | 0.46 | 0.51 |
| Shrub/ Scrub | 332 | 67208 | 1620 | 240 | 0.97 | 0.58 | 0.98 | 0.17 | 0.39 |
| Grassland/ Herbaceous | 331 | 66787 | 1765 | 517 | 0.97 | 0.39 | 0.97 | 0.16 | 0.3 |
| Pasture/ Hay | 1955 | 62555 | 2812 | 2078 | 0.93 | 0.48 | 0.96 | 0.41 | 0.47 |
| Cultivated Crops | 20135 | 42113 | 553 | 6599 | 0.90 | 0.75 | 0.99 | 0.97 | 0.79 |
| Woody Wetlands | 5580 | 57257 | 1323 | 5240 | 0.91 | 0.52 | 0.98 | 0.81 | 0.56 |
| Emergent Herbaceous Wetlands | 3547 | 61081 | 2570 | 2202 | 0.93 | 0.62 | 0.96 | 0.58 | 0.61 |

Table S4. Area of leafy spurge invasion in square kilometers predicted from the TempCNN model between 2000 and 2020.

| Years | Invaded Area (sq. km) |
|---|---|
| 2000 - 2002 | 1,067 |
| 2003 - 2005 | 2,417 |
| 2006 - 2008 | 4,149 |
| 2009 - 2011 | 3,124 |
| 2012 - 2014 | 1,372 |
| 2015 - 2017 | 2,134 |
| 2018 - 2020 | 7,156 |

Table S5. Sequential analysis of deviance table for quasipoisson generalized linear model testing for a relationship between the predicted extent of leafy spurge invasion with mean Palmer Drought Severity Index (PDSI) values and across different geographic regions. Significant P-values (*P < 0.05*) are shown in bold.

| Source | df | $X^2$ | F | Deviance | Resid. Deviance | P |
|---|---|---|---|---|---|---|
| NULL | 62 | | | | 44002571 | |
| Mean PDSI | 1 | 50.04 | 35.87 | 4496364 | 39506206 | **< 0.001** |
| Region | 8 | 266.93 | 33.37 | 33458118 | 6048088 | **< 0.001** |
| Mean PDSI * Region | 8 | 3.90 | 0.49 | 487446 | 5560642 | 0.867($X^2$) / 0.860 (F) |
| Residual | 45 | | | | | |

*Dispersion parameter = 125344
